# A transporter essential for apical secretory organelles homeostasis in *Toxoplasma gondii*

**DOI:** 10.64898/2026.10.23.760958

**Authors:** Alessandro Bonavoglia, Rachel Humann, Oscar Vadas, Isabel Meister, Bohumil Maco, David J Dubois, Dominique Soldati-Favre, Syrian G Sanchez

**Author notes:** Correspondence: Syrian G Sanchez; and Dominique Soldati-Favre.

## Abstract

Apicomplexan parasites rely on the rhoptries and micronemes to coordinate egress, motility, and host cell invasion. In *Toxoplasma gondii*, mechanistic insights into the control of these organelles’ lipid composition remain sparse. Although nearly 60 *T. gondii* Major Facilitator Superfamily (MFS) are predicted, many remain functionally uncharacterized. The MFS protein HAP12 was previously annotated as an apicoplast protein contributing to invasion and organelle inheritance. Based on localization and phylogenetic analyses, we show that HAP12 (renamed TFP5), is predominantly associated with the Golgi, with additional signal in the ELC and apical compartments, without convincing evidence of a major apicoplast-associated pool. Furthermore, TFP5 clusters with a subgroup of MFS transporter facilitator proteins (TFPs) involved in the morphological integrity of secretory apical organelles. TFP5 depletion results in defective microneme secretion and pronounced alterations in rhoptry morphology and positioning. Structural analyses revealed that TFP5 is strongly similar to MFSD2-like lipid transporters, while lipidomic analyses showed substantial changes in parasite lipid composition following TFP5 depletion. Together, these findings support a model in which TFP5 regulates lipid homeostasis required for the function of apical secretory organelles, thereby critically contributing to parasite virulence.

**IMPORTANCE:** The Apicomplexa relies on tightly regulated secretion from specialized apical organelles for host cell invasion and intracellular proliferation, processes essential for pathogenicity. While the membrane trafficking pathways controlling these organelles are well characterized in *Toxoplasma gondii*, the contribution of membrane lipid transporters remains poorly understood. Here, we identify TFP5 as an Alveolata-conserved MFSD2-like transporter localized to the endomembrane system and micronemes. Consistent with a role in membrane lipid homeostasis, TFP5 depletion alters the parasite lipidome, severely impairs microneme secretion and rhoptry morphology, and consequently disrupts motility, egress, and host cell invasion without affecting intracellular replication. These findings reveal a critical role for membrane lipid homeostasis in the organization and function of apical secretory organelles and identify TFP5 as a potential therapeutic target against apicomplexan parasites.

## INTRODUCTION

Apicomplexan parasites, including *Toxoplasma gondii*, *Plasmodium* spp., and *Cryptosporidium* spp., are major human and animal pathogens (1–3). Their success as obligate intracellular parasites relies on coordinated host-cell invasion, intracellular replication, and egress (4). Two apical secretory organelles, the micronemes and rhoptries, are central to these processes (5). In *T. gondii*, micronemes secrete perforins, adhesins, and motility factors required for egress, gliding, and invasion, whereas rhoptries deliver neck (RON) and bulb (ROP) proteins that mediate moving junction formation and remodel host cell functions (6–8).

Micronemes and rhoptries are assembled de novo during daughter cell formation through a specialized branch of the endosomal trafficking pathway (9). Newly synthesized precursor proteins exit the endoplasmic reticulum, transit through the Golgi apparatus, and are sorted through post-Golgi compartments, including the endosomal-like compartment (ELC) and plant-like vacuole (PLVAC), before reaching the apical secretory organelles. (6, 7, 10). Consistent with this model, disruption of proteins associated with these compartments impairs secretory organelle biogenesis or cargo trafficking (11–21).

Besides correct trafficking, many proteins destined for the apical secretory organelles require proteolytic maturation to become functional. This occurs along the endosomal pathway, where the endosomal-like compartment (ELC) serves as the main sorting and maturation hub and the acidic plant-like vacuole (PLVAC) contributes to protein processing (22, 23). These compartments are functionally interconnected, as illustrated by the PLVAC-resident protease cathepsin L (CPL), which is also detected in the ELC and is required for microneme maturation (23). Another key component is the ELC-resident aspartyl protease 3 (ASP3), which processes multiple microneme and rhoptry proteins and is essential for parasite egress and invasion. Although ASP3 depletion does not affect microneme trafficking, it blocks proteolytic maturation, disrupts rhoptry neck morphology, and prevents rhoptry discharge (25, 26). Membrane lipid composition is a fundamental determinant of vesicle formation, membrane curvature, and fusion (27). Apical secretory organelles are highly specialized in apicomplexan parasites, implying unique molecular composition. While their protein content has been extensively characterized, their lipid composition remains largely unknown because these organelles are difficult to isolate (28–30).

Lipid transfer proteins and membrane transporters are thus expected to regulate lipid homeostasis along the trafficking pathways and within the organelles. Among them, Major Facilitator Superfamily (MFS) proteins form a conserved class of transporters that mediate the movement of diverse substrates across membranes (31). While most MFS proteins transport metabolites, some atypical MFS domain-containing proteins (MFSDs), as well as MFS Spinster proteins, facilitate lipid transport (32–36). In *T. gondii*, several MFS transporters localize to specific subcellular compartments and participate in processes including metabolism, organelle biogenesis, and exocytosis (37–46). A subset of these transporters, termed Transporter Facilitator Proteins (TFPs), is conserved among Apicomplexa and localizes to the Golgi apparatus, the ELC, or apical secretory organelles, where some members contribute to the biogenesis and maturation of these specialized compartments (40).

The MFS transporter TGGT1_226020, previously annotated as hypothetical apicoplast transporter 12 (HAP12) (37, 43), was initially linked to defects in host cell invasion and a delayed death phenotype (47). Here, we show that HAP12, renamed TgTFP5, is an Alveolata-conserved MFS transporter that clusters with other TgTFPs and is structurally related to the *Danio rerio* lysophosphatidylcholine (LPC) transporter MFSD2A (48). Our results shows that TgTFP5 predominantly localizes to the Golgi and partially colocalize to the ELC, and micronemes. Its conditional depletion impairs microneme secretion without affecting organelle morphology while severely disrupting rhoptry architecture. These findings underscore the importance of secretory organelle homeostasis for parasite virulence.

## RESULTS

### TFP5 is an MFS transporter conserved across most Alveolata

From a previous screen of fitness-conferring MFS transporters (46), we selected TGGT1_226020 for further characterization based on its low fitness score (49) and predicted micronemal localization by LOPIT (50). TGGT1_226020 encodes a 12-transmembrane-domain protein, as identified by DeepTMHMM, and a conserved MFS transporter domain detected by InterPro (**Fig. 1A**). BLAST analysis identified TFP5 homologues across several coccidian and alveolate lineages, but not in piroplasmids or *Cryptosporidium* species (**Fig. S1A** and **B**; **Table S1** for BlastP scores and E-values). AlphaFold predictions for TGGT1_226020 (**Fig. 1B**) and PF3D7_0529200 (**Fig. S1C**) showed a strong structural alignment (**Fig. S1D**), supporting an evolutionary relationship between the two proteins. Phylogenetic analysis using the previously generated dataset (46) placed TGGT1_226020 within the *T. gondii* TFP transporter family (**Fig. 1C**), leading us to designate it as TgTFP5. These findings support the evolutionary conservation of TFP5 in Apicomplexa and Colpodellida lineages, suggesting preservation of its biological function.

**Fig. 1.**
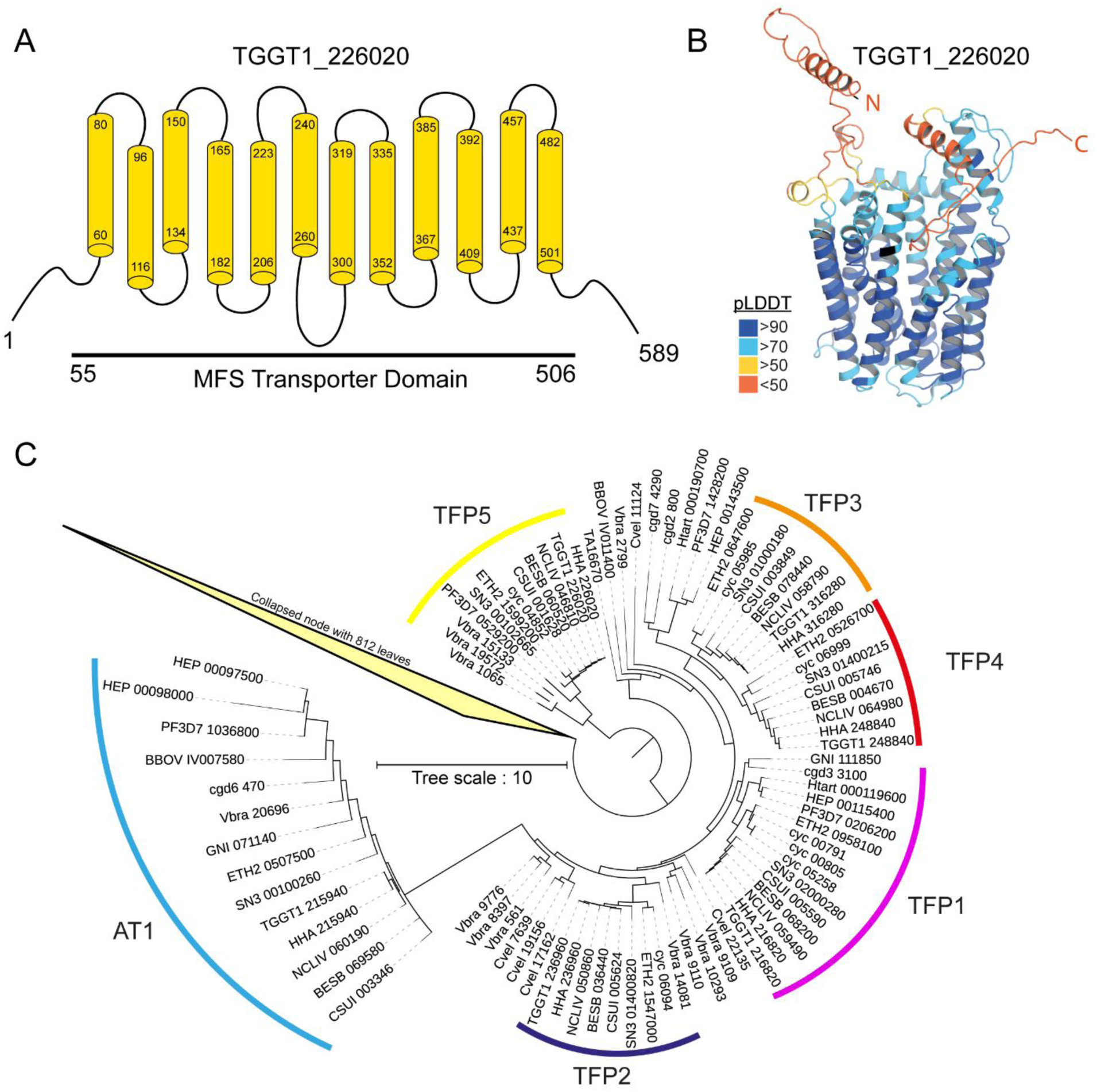
Structural and phylogenetic analysis of the *T. gondii* MFS transporter TGGT1_226020 (TFP5). **(A)** Schematic representation of the TGGT1_226020 putative MFS transporter. Transmembrane domains (TMs) are shown in yellow. Amino acid numbers indicate positions along the full-length protein, the start and end of each TM domain as predicted by TMHMM, and the boundaries of the predicted MFS transporter domain identified by InterPro. **(B)** AlphaFold3 prediction of the 3D structure of TGGT1_226020, coloured according to the pLDDT score. The first N-terminal (N) and last C-terminal (C) residues of the predicted protein-coding sequences are annotated on the structure. **(C)** Collapsed phylogenetic tree extracted from the previously published phylogeny of MFS transporters across different Alveolata species (46). The tree was reconstructed from homologous protein sequences identified in Apicomplexa and related lineages (VEuPathDB gene IDs indicated). Five distinct clades corresponding to the TFP1–TFP5 subfamilies are highlighted with colored outlines (TFP1 in pink, TFP2 in dark blue, TFP3 in red, TFP4 in light red/brown, and TFP5 in yellow). The AT1 group (light blue) represents a divergent but related MFS subfamily. A collapsed node containing 812 sequences is shown to simplify tree visualization. Branch lengths reflect evolutionary distances (scale bar = 10 substitutions per site).

### TFP5 is an evolutionarily conserved MFSD2-like transporter with structural hallmarks of lipid binding

Phylogenetic analysis including human MFS transporters, TgTFP5 and representative apicomplexan homologues placed TFP5 proteins in a distinct clade closely related to the lipid transporters MFSD2A and MFSD2B, identifying TFP5 as an MFSD2-like lineage within the MFS superfamily (**Fig. S1E**). Consistent with this relationship, structural similarity analysis using FoldSeek (51) identified *Danio rerio* MFSD2A (PDB: 8D2S) (48), an Na-dependent LPC symporter, as the closest structural homologue. Overlay of TgTFP5 or PfTFP5 MFS domains with DrMFSD2A’s cryo-EM structure revealed extensive structural conservation (**Fig. 2A to C**). Comparison of electrostatic **(Fig. 2D to I)** and hydrophobic **(Fig. 2J to R)** properties within the central cavities of TgTFP5, PfTFP5, and DrMFSD2A revealed highly similar physicochemical environments, consistent with a potential role in the recognition or transport of LPC or related lipids.

**Fig. 2.**
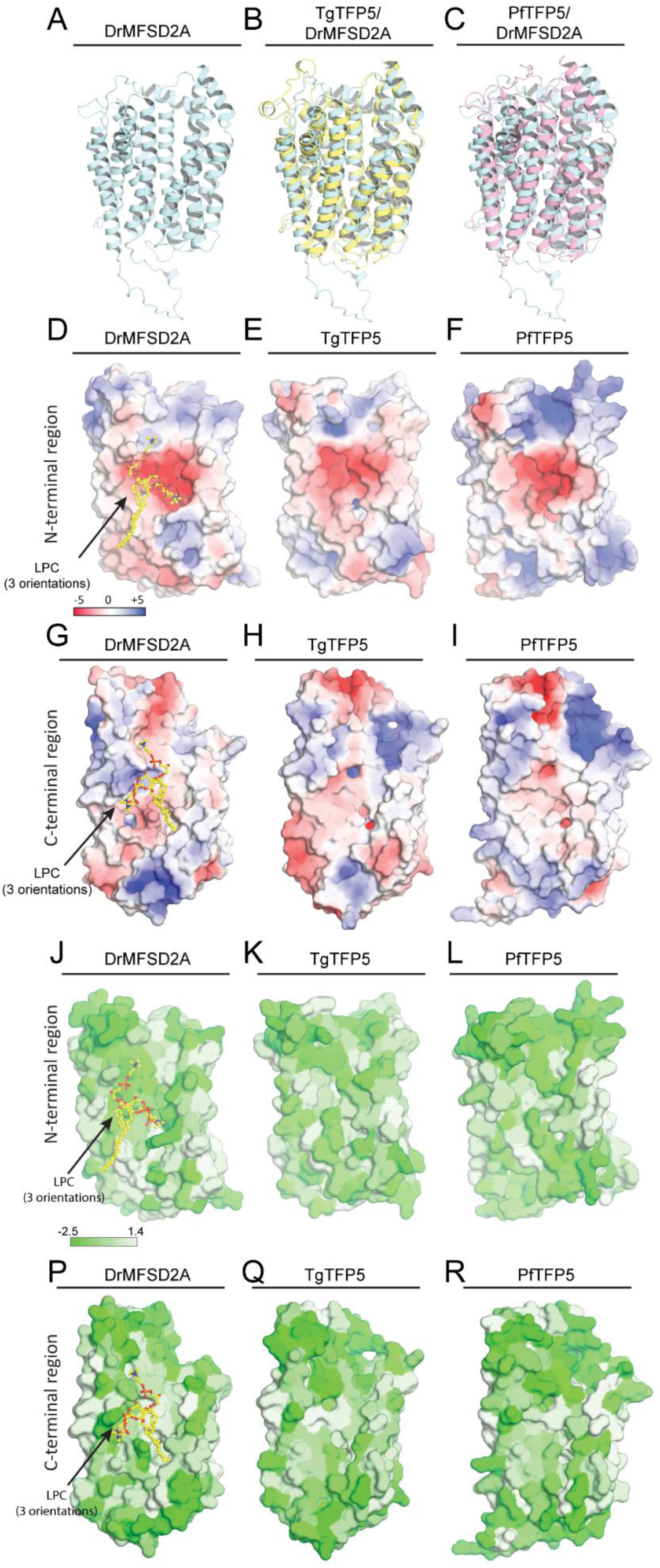
DrMFSD2A, TgTFP5 and PfTFP5 share overall structure and binding pocket electrostatic and hydrophobicity properties. **(A)** Cryo-EM structure of DrMFSD2A in the inward-open conformation (PDB: 8D2S). Structural alignment of **(B)** TgTFP5 and **(C)** PfTFP5 high-confidence regions from AlphaFold models with the DrMFSD2A cryo-EM structure. **(D-I)** Surface representations of the internal cavity of the N-terminal half **(D-F)** or C-terminal half **(G-I)** of the DrMFSD2A **(D, G)**, TgTFP5 **(E, H)**, and PfTFP5 **(F, I)** transporters in the same orientation, coloured according to electrostatic potential. Electrostatic potential ranges from −5 kT/e (red, negative) to +5 kT/e (blue, positive). **(J–R)** Surface representations of the internal cavity of the N-terminal half **(J–L)** or C-terminal half **(P– R)** of the DrMFSD2A **(J, P)**, TgTFP5 **(K, Q)**, and PfTFP5 **(L, R)** transporters in the same orientation, coloured according to hydrophobicity (Eisenberg scale), ranging from highly hydrophobic (white) to highly hydrophilic (green). LPC orientations are shown for DrMFSD2A structures bound to the lipid in three orientations (PDBs: 8D2S, 8D2U, 8D2V).

### TFP5 localizes to the early secretory, endosomal-like compartment and micronemes

To address the localization and function of TFP5, we generated a conditional knockdown line using the U1 silencing system in the DiCre ΔKu80 ΔHXGPRT background (**Fig. S2A and B).** Western blot analysis confirmed expression of TFP5-3Ty and revealed multiple protein isoforms (**Fig. S2C**). We next investigated its localization by immunofluorescence assays (IFA) using an anti-Ty antibody and markers of various subcellular compartments. TFP5 displayed a perinuclear distribution, consistent with localization along the secretory pathway and an apical gradient reminiscent of micronemal proteins (**Fig. 3A**).

**Fig. 3.**
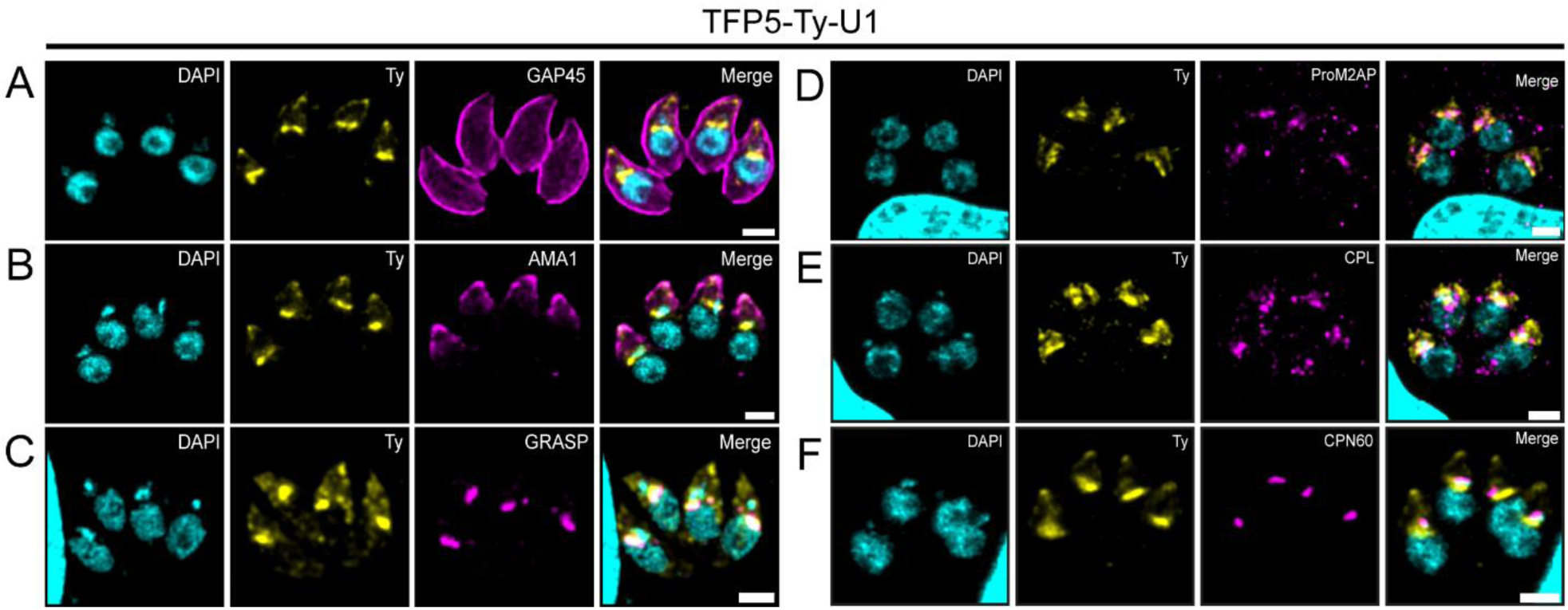
Subcellular localization of TFP5 in *T. gondii*. **(A-F)** Colocalization analysis of TFP5 (yellow) with various *T. gondii* markers (magenta). **(A)** Overview of TFP5 localization using co-staining with the IMC marker GAP45. **(B)** TFP5-3Ty shows partial overlap with the microneme marker AMA1 at the parasite apex. **(C)** Parasites expressing the Golgi marker GRASP-YFP show strong colocalization with TFP5. **(D)** The ELC marker ProM2AP shows partial colocalization with TFP5. **(E)** The PLVAC marker CPL shows partial colocalization with TFP5. **(F)** TFP5 shows no convincing colocalization with the apicoplast marker CPN60. Scale bars: 2 μm.

These observations were confirmed by the partial colocalization of TFP5-3Ty with the micronemal marker AMA1 **(Fig. 3B)**. We next assessed co-localization with secretory pathway markers, including GRASP55-YFP for the cis-Golgi (52), proM2AP for the ELC (23), and Cathepsin-L for the PLVAC (53)(**Fig. 3C to E**). TFP5-3Ty showed strong co-localization with Golgi and partial overlap with ELC. However, TFP5 showed limited overlap with CPL. Co-staining with the apicoplast marker CPN60 did not reveal convincing colocalization with TFP5-3Ty, although the presence of a minor apicoplast-associated pool cannot be formally excluded (**Fig. 3F)**. Additionally, solubility assays confirmed that TFP5 displays the biochemical properties of an integral membrane protein **(Fig. S2D).** In summary, these observations indicate that TFP5-3Ty is an integral membrane protein predominantly associated with the endomembrane system, with additional signal detected in apical compartments partially overlapping with micronemes.

### TFP5 is required for progression through the *T. gondii* lytic cycle

To investigate the role of TFP5 during the *T. gondii* lytic cycle, we first used the TFP5-Ty-U1 strain, which recapitulated the expected localization pattern. Rapamycin treatment efficiently depleted TFP5 within 72 h (**Fig. S2B, C and E**), resulting in a severe parasite propagation defect (**Fig. S3A and B**). To establish a faster depletion system, we generated a TFP5-mAID-HA **(Fig. S4A and B)**. TFP5 was efficiently depleted within one hour of auxin (IAA) treatment **(Fig. S4C)**. Although the TFP5-mAID-3HA predominantly colocalized with the golgi,marker GRASP55 suggesting altered trafficking of the mAID-3HA-tagged protein, IAA treatment confirmed efficient protein degradation, supporting the cytosolic accessibility of the C-terminal AID tag (**Fig. S4D and E**). However, untreated TFP5-mAID-HA parasites formed smaller plaques compared to the TIR1-IAA condition, suggesting that the tagged parasite fitness is partially compromised. Nonetheless, Plaque assays revealed a marked defect following TFP5 depletion **(Fig. 4A and B)**. Together, these two independent depletion systems establish TFP5 as essential for efficient parasite propagation.

**Fig. 4.**
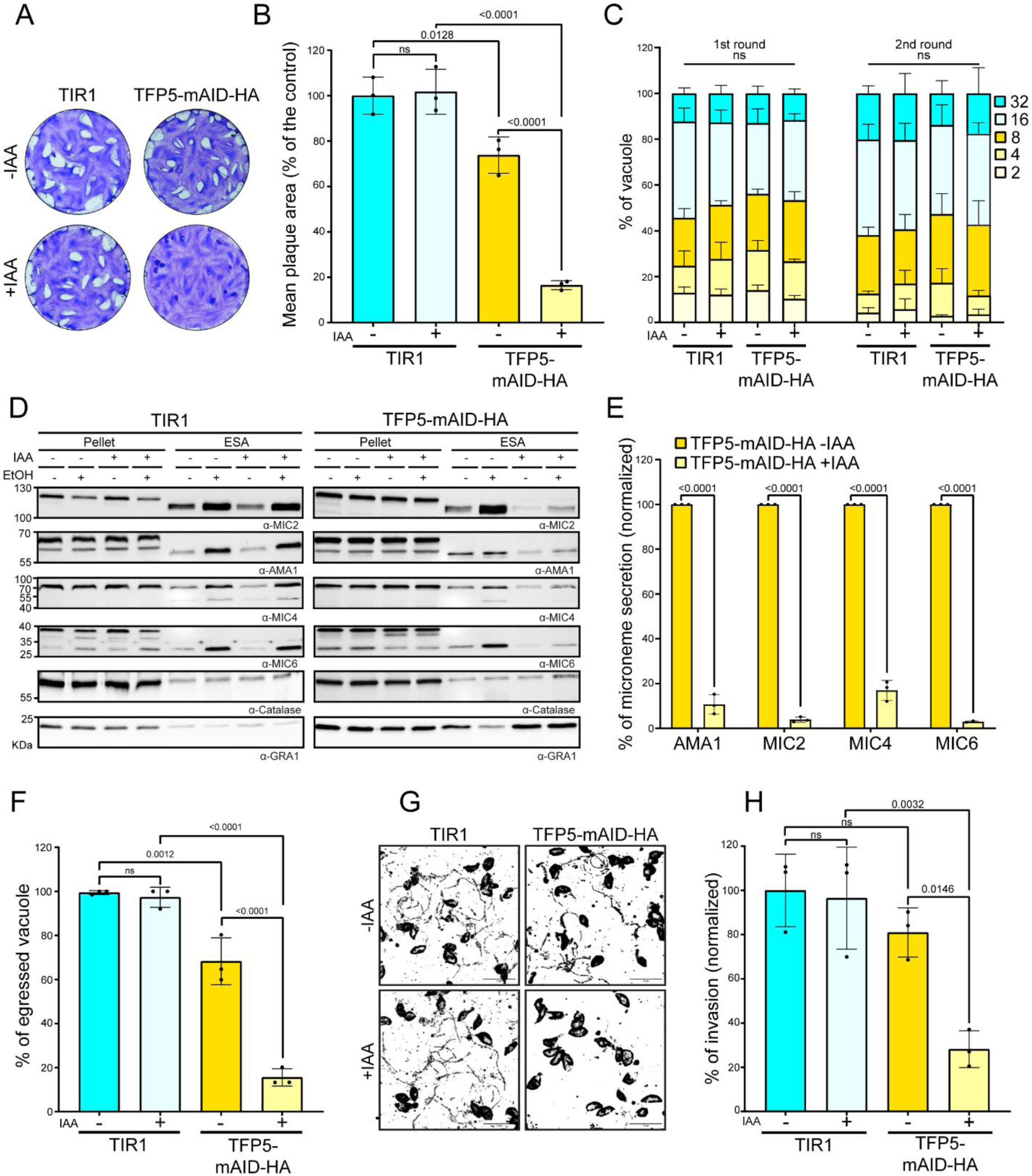
TFP5 is required for efficient microneme secretion and parasite lytic cycle progression. **(A)** Representative images from plaque assays performed in three independent experiments showing lytic cycle progression in parental TIR1 and TFP5-mAID-HA parasites, with or without IAA treatment. **(B)** Quantification of plaque assay results for TIR1 and TFP5-mAID-HA parasites ± IAA. Plaque areas were normalized to the TIR1-IAA condition. Statistical analysis was performed using one-way ANOVA followed by Tukey’s multiple comparisons test (mean ± SD; n = 3 biologically independent replicates). **(C)** Quantification of parasite numbers per vacuole during the first and second rounds of intracellular replication in parental TIR1 and TFP5-mAID-HAstrains ± IAA. Statistical differences between strains within each category were analyzed using two-way ANOVA with Tukey’s multiple comparisons test (mean ± SD; n = 3 biologically independent replicates). **(D)** Immunoblot analysis of microneme secretion in TIR1 and TFP5-mAID-HA parasites treated ± IAA after ethanol (EtOH) stimulation. Secretion of MIC2, AMA1, MIC4, and MIC6 was assessed by analysing their distribution in pellet (P) and supernatant (S) fractions. Anti-GRA1 was used as a control for constitutive secretion, and catalase served as a control for parasite lysis. Representative images of three biological replicates. **(E)** Quantification of AMA1, MIC2, MIC4, and MIC6 secretion into the supernatant following EtOH stimulation. Secretion levels were first normalized to constitutive GRA1 secretion for each sample and then expressed relative to the TFP5-mAID-HA-IAA condition. Statistical significance was assessed using two-way ANOVA with Tukey’s multiple comparisons test (mean ± SD; n = 3 biologically independent replicates). **(F)** Quantification of parasite egress in TIR1 and TFP5-mAID-HA parasites ± IAA following stimulation with the egress-inducing compound BIPPO. Statistical significance was determined using one-way ANOVA followed by Tukey’s multiple comparisons test (mean ± SD; n = 3 biologically independent replicates). **(G)** Representative images of parasite gliding after BIPPO stimulation in TIR1 and TFP5-mAID-HA parasites treated ± IAA for 30 hours. Scale bar: 10 μm. **(H)** Quantification of intracellular TIR1 and TFP5-mAID-HA parasites after 30 h of pretreatment ± IAA. Invasion efficiency was measured 30 minutes post-infection and expressed relative to the TIR1 −IAA control. Statistical significance was assessed using one-way ANOVA followed by Tukey’s multiple comparisons test (mean ± SD; n = 3 biologically independent replicates).

### TFP5 is essential for microneme secretion and microneme-dependent processes

We next examined TFP5’s contribution to different steps of the lytic cycle. TFP5 depletion did not affect intracellular replication over one or two lytic cycles, excluding a delayed-death phenotype **(Fig. 4C).** In contrast, ethanol-induced microneme secretion was strongly impaired in both iKD systems after TFP5 depletion (**Fig. 4D and 4E and S3C and D**). Consistent with this defect, TFP5-depleted parasites displayed markedly reduced BIPPO-induced egress (54), gliding motility, and host cell invasion **(Fig. 4F-H and S3E and F)**. Although untreated TFP5-mAID-HA parasites exhibited mild basal egress and invasion defects, both depletion systems produced comparable phenotypes. Overall, these findings establish TFP5 as essential for microneme secretion and microneme-dependent processes, including egress, gliding motility, presumably rhoptry discharge and invasion.

### Loss of TFP5 disrupts rhoptry morphogenesis and microneme collar organization but not protein trafficking or processing

Because TFP5 localizes to compartments involved in secretory organelle biogenesis, we first examined the localization of representative microneme proteins. AMA1, MIC2, MIC3, and MIC4 retained their characteristic apical localization in both TFP5 depletion systems, indicating that microneme protein trafficking is largely preserved (**Fig. 5A-D and S5A-D**)

**Fig. 5.**
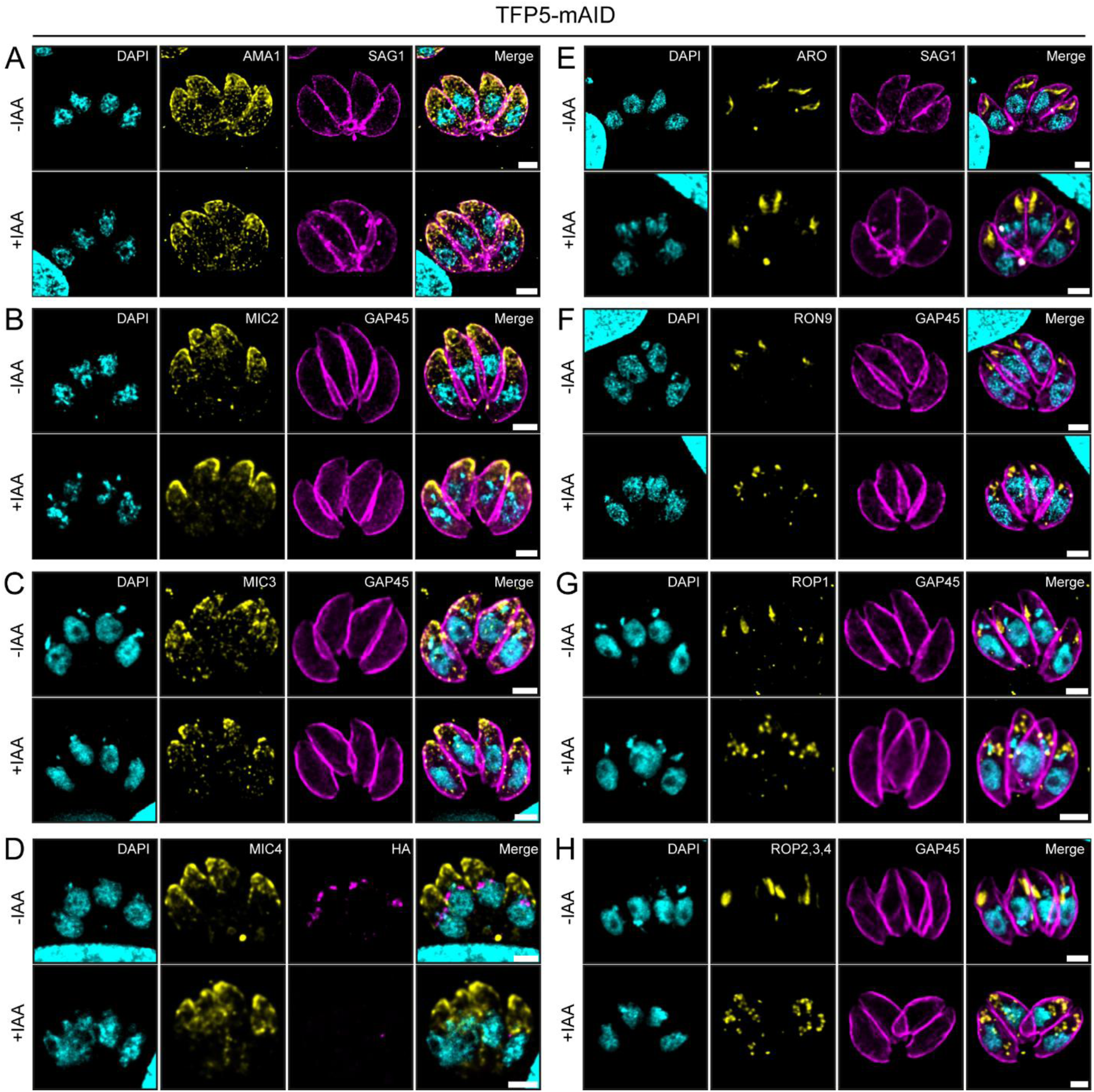
TFP5 depletion does not alter the overall microneme protein trafficking but disrupts normal rhoptry morphology. **(A–D).** The microneme staining pattern is not affected by TFP5 depletion. **(A)** Anti-AMA1, **(B)** Anti-MIC2, **(C)** Anti-MIC3 and **(D)** Anti-MIC4 (yellow) were co-stained with anti-SAG1 to label the plasma membrane, anti-GAP45 to label the parasite IMC or anti-HA to highlight the TFP5-mAID-HA protein (magenta). **(E–H)** Rhoptry morphology is altered upon TFP5 depletion. **(E)** Anti-ARO **(F)** Anti-RON9, **(G)** Anti-ROP1, and **(H)** Anti-ROP2/3/4 (yellow) were co-stained with anti-SAG1 or anti-GAP45(magenta). DAPI marks the nucleus in all IFAs. Scale bars: 2 µm.

To examine whether rhoptry protein trafficking was altered upon TFP5, we stained ARO, RON9, ROP1, and ROP2–4 in both iKD mutants (**Fig. 5E to H and S5E to H**). ARO staining remained associated with rhoptries in TFP5-depleted parasites, although rhoptry morphology appeared markedly altered, adopting a more compact and less elongated appearance compared with controls. RON9, ROP1, and ROP2–4 displayed a more vesicular staining pattern that lacked the characteristic elongated morphology of mature rhoptries upon TFP5 depletion. Transmission electron microscopy (TEM) of TFP5-depleted parasites revealed unchanged microneme abundance but highlighted a defect in microneme collar organization (**Fig. 6A**). Moreover, the typical club-shaped architecture of rhoptries was profoundly disrupted, with shortened necks and enlarged bulbs, resulting in rounder and more distorted organelles. In addition, some rhoptries failed to properly cluster at the parasite apex and appeared misoriented within the cytoplasm (**Fig. 6B**). To further characterize the rhoptry morphological defects observed following TFP5 depletion, we quantified the proportion of parasites displaying normal rhoptry morphology by transmission electron microscopy. Among the 43 parasites examined under control conditions (−IAA), approximately 97% exhibited normal rhoptry morphology. In contrast, only approximately 9% of the 36 parasites analyzed following auxin treatment (+IAA) displayed normal rhoptry morphology (**Fig. S6A**).

**Fig. 6.**
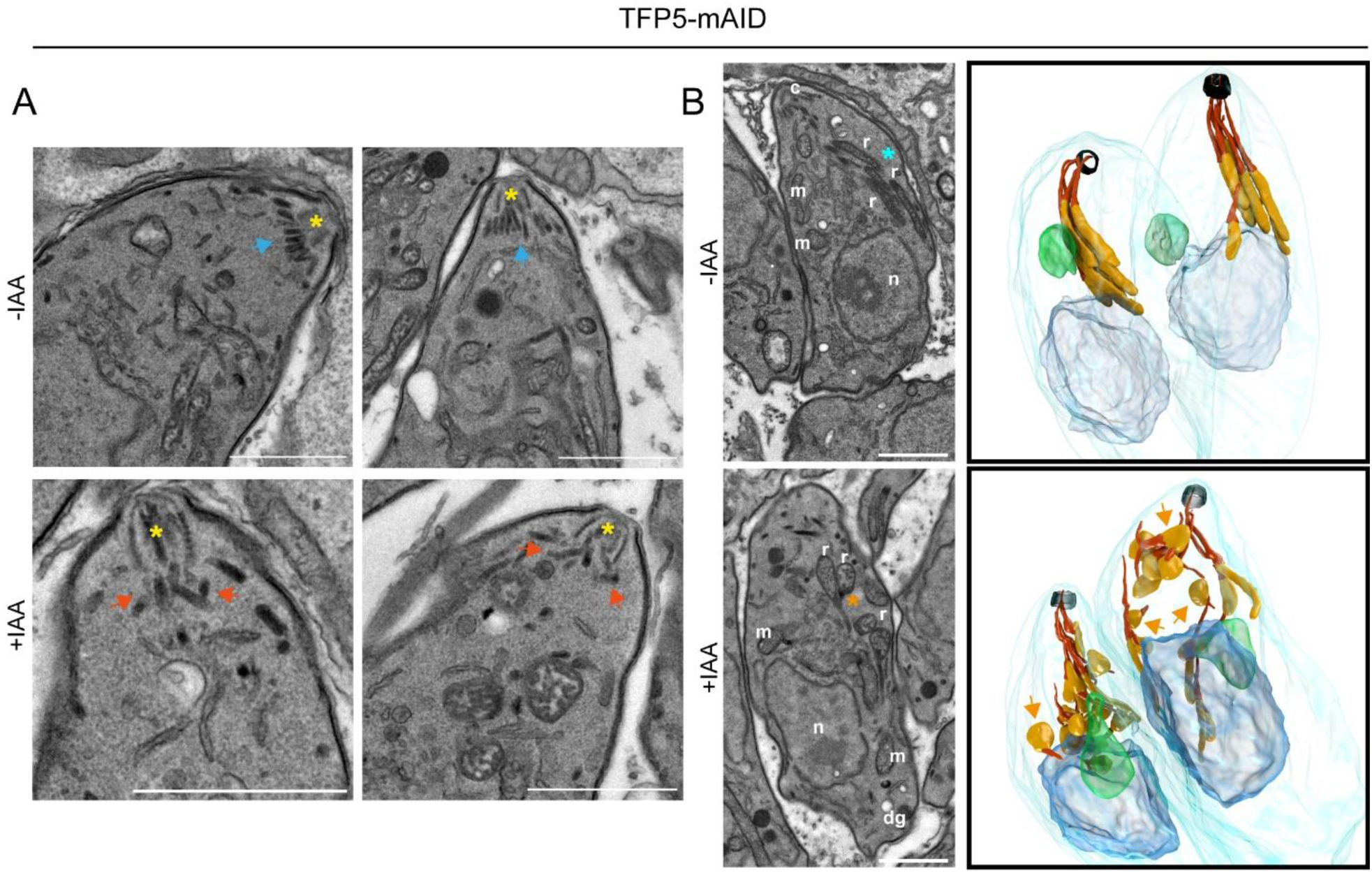
Ultrastructural analysis reveals altered collar microneme organization and rhoptry morphology upon TFP5 loss. **(A)** Transmission electron microscopy of TFP5-mAID-HA parasites grown in the presence or absence of IAA shows that under control conditions (–IAA), apical micronemes are organized in a characteristic collar-like arrangement (Blue arrows) beneath the conoid (yellow asterisk). Upon IAA treatment, micronemes appear dispersed and disorganized within the apical region (dark orange arrows). Scale bars: 1 µm. **(B)** Electron microscopy analysis of TFP5-depleted parasites reveals pronounced defects in rhoptry morphology and positioning. Representative EM section of -IAA or +IAA-treated parasites is shown on the left. Cyan asterisks highlight the presence of normal rhoptries, while light orange asterisks indicate the presence of morphologically abnormal rhoptries. Scale bar: 1 µm. Another Focused Ion Beam Scanning Electron Microscopy (FIBSEM) reconstruction is displayed on the right. In the absence of TFP5, the rhoptry bulb (light orange) appears enlarged, while the rhoptry neck (dark orange) is shortened compared to the control condition. In addition, light orange arrows highlight impaired rhoptry clustering. The parasite plasma membrane is highlighted in light blue. The nuclei are coloured in dark blue, and the apicoplast is highlighted in green.

The discrepancy between the lack of detectable TFP5 localization at the rhoptry and the pronounced morphological phenotype of this organelle upon TFP5 depletion prompted us to examine whether the Golgi and post-Golgi compartments were affected. The localization of GRASP, CPL, and proM2AP markers showed a normal staining pattern in the absence of TFP5 (**Fig. S6B to D**). We also examined the effect of TFP5 depletion on dense granules, the apicoplast, and the mitochondrion using GRA1, CPN60, and F1-ATPase as markers, respectively. No alteration of staining or morphological abnormalities were detected for these organelles (**Fig. S6E to G**). Likewise, proteolytic maturation of representative microneme and rhoptry proteins remained unaffected following TFP5 depletion compared to the ASP3ty-IKD control strain (25) (**Fig. S6H**), indicating that TFP5 is dispensable for cargo processing. Together, these findings support a role for TFP5 in the structural organization of apical secretory organelles rather than protein trafficking and maturation

### Global lipid profiling reveals alterations in parasite lipid homeostasis upon TFP5 depletion

To determine whether the structural defects observed upon TFP5 depletion were associated with altered membrane lipid homeostasis, we performed untargeted lipidomics. In total, 942 lipid species were identified, and OPLS-DA revealed clear separation between control and TFP-depleted parasites, indicating extensive lipid remodelling (**Fig. S7A to C**).

To specifically assess the lipidomic changes associated with TFP5 depletion, we calculated fold changes in lipid abundance between auxin-treated and untreated TFP5-mAID-HA parasites using paired biological replicates (**Fig. 7**). The corresponding lipid abundances, normalized to the parental TIR1 +IAA control, are also presented to distinguish depletion-associated changes from alterations already present in untreated TFP5-mAID-HA parasites (**Fig. S8**).

**Figure 7.**
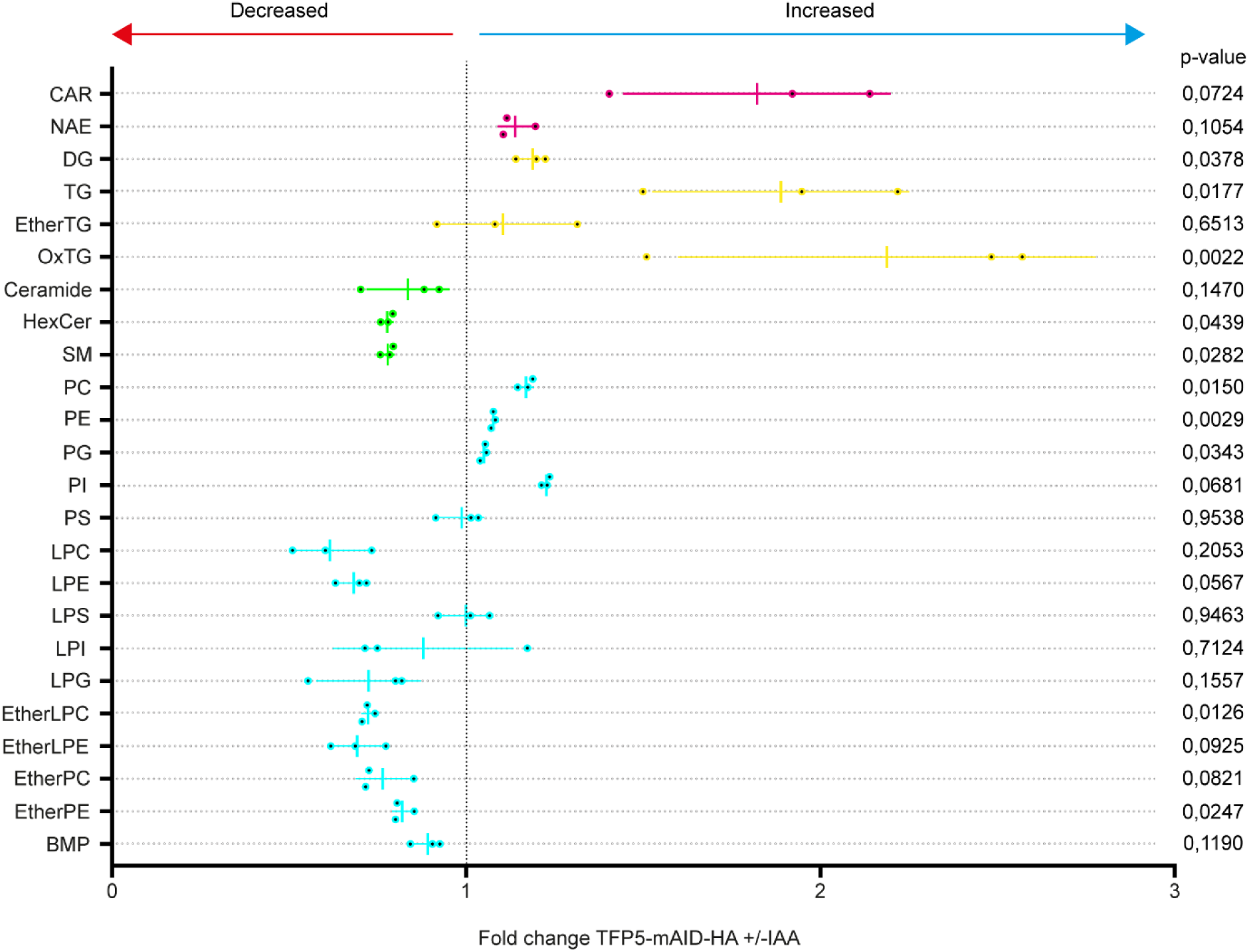
Lipidomic alterations associated with TFP5 depletion. Dot plot showing fold changes in the indicated lipid classes in TFP5-mAID-HA parasites following auxin treatment (+IAA) relative to untreated parasites (−IAA). Fold changes were calculated for each biological replicate by dividing the normalized lipid abundance in the +IAA condition by that of the corresponding −IAA condition from the same independent experiment. The vertical dashed line indicates a fold change of 1 (no change), with values below and above 1 representing decreased and increased lipid abundance, respectively. Lipid categories are color-coded as follows: fatty acyls (magenta), glycerolipids (yellow), sphingolipids (green), and glycerophospholipids (cyan). Each dot represents one paired biological replicate (n = 3), and bars indicate the mean ± SD. Statistical comparisons between TFP5-mAID-HA +IAA and −IAA were performed using a one-way repeated-measures ANOVA followed by Tukey’s multiple comparisons test, including the parental TIR1 +IAA condition in the analysis. Adjusted p-values for the indicated comparison are shown on the right. The corresponding normalized lipid abundances for all three experimental conditions are presented in **Fig. S8**.

At the lipid class level, TFP5 depletion was associated with changes across several lipid categories, although the magnitude and direction of these changes varied between classes (**Fig. 7**). Among fatty acyls, acylcarnitines increased following TFP5 depletion, whereas N-acylethanolamines remained largely unchanged (**Fig. 7 and S8A**). Glycerolipids were also affected, with increased diacylglycerol (DG) and triacylglycerol (TG) levels following auxin treatment. Oxidized triacylglycerols (OxTG) were similarly enriched, whereas ether-linked TG levels remained largely unchanged (Fig. 7 and S8B). Notably, DG levels were already elevated in untreated TFP5-mAID-HA parasites relative to the parental TIR1 control, indicating that part of the lipid remodelling preceded auxin-induced depletion.

Among sphingolipids, ceramide levels remained largely unchanged, whereas hexosylceramides (HexCer) and sphingomyelins (SM) decreased following TFP5 depletion (**Fig. 7 and S8C**). Glycerophospholipids also showed substantial remodelling, with increased phosphatidylcholine (PC) and phosphatidylethanolamine (PE) levels following IAA treatment. In contrast, phosphatidylglycerol (PG), phosphatidylinositol (PI), and phosphatidylserine (PS) exhibited variable trends without consistent significant changes. Lysophospholipids, including LPC, LPE, LPS, LPI, and LPG, generally showed reduced abundance in the mutant background, although these differences did not consistently reach statistical significance. Ether-linked lysophospholipids, including etherLPC and etherLPE, decreased following TFP5 depletion, while etherPE was also reduced. Finally, bis(monoacylglycero)phosphate (BMP) was elevated in the mutant background, including in untreated parasites (**Fig. 7 and S8D**).

Together, these analyses reveal lipid remodelling associated with TFP5 depletion, characterized by increased abundance of several glycerolipid and glycerophospholipid classes and reduced abundance of selected sphingolipids. The complementary comparison with the parental TIR1 control further indicates that some lipid alterations are already present in untreated TFP5-mAID-HA parasites, while others become more pronounced following TFP5 depletion. These findings support a role for TFP5 in maintaining parasite lipid homeostasis.

## DISCUSSION

MFSs represent one of the most expanded groups of membrane transporters across all domains of life. In *T. gondii*, close to 60 proteins are predicted to contain an MFS domain, highlighting the prominence of this family within the parasite’s transportome (55). This abundance likely reflects the need for regulated transport of small solutes between extracellular and intracellular environments and for exchanges between intracellular organelles. Consistent with this idea, genome-wide functional screens have identified several MFS transporters as strong contributors of tachyzoite fitness, underscoring their central role in cellular homeostasis and survival in this stage (49).

This study establishes a critical role for TFP5 in controlling secretory organelles’ morphology and function, ultimately important for parasite fitness. Until recently, the reported role of this protein in invasion, together with its apicoplast localization, remained confusing (37, 43). The 6Ty-tagged protein showed a dual localization within the secretory pathway and an apical gradient consistent with micronemal staining (37), in agreement with LOPIT analyses (50). In a subsequent study using a large AID-6Ty fusion protein, TFP5 was reported to localize to the apicoplast (43). These discrepancies suggested that the large AID degron alters TFP5 localization. Here, using a shorter 3Ty tag, we observed a predominantly Golgi-associated distribution of TFP5, with additional signal in the ELC and apical compartments partially overlapping with micronemes, consistent with the first report (37) and placing it at a key position to be involved in the biogenesis of apical secretory organelles. Although neither tagged version showed convincing colocalization with the apicoplast marker CPN60, our imaging data cannot formally exclude a minor apicoplast-associated pool of TFP5.

Phylogenetic analysis revealed that TFP5 clusters within the TFP transporter family, whose members exhibit distinct subcellular localizations, ranging from the Golgi apparatus to the ELC and secretory organelles. TFP5 is conserved in Coccidia and *Laverania* lineages but is absent from rodent malaria parasites and other apicomplexan lineages. Such a restricted conservation of TFP5 suggests that this transporter may contribute to specialized features of the apical secretory system in specific alveolate lineages. Consistent with a conserved function in *Laverania*, although PfTFP5 has not been experimentally localized, genome-wide fitness screens identified it as an important determinant of asexual blood-stage growth (56).

Using two independent conditional knockdown systems, we demonstrate that TFP5 is essential for parasite fitness. The DiCre U1-mediated knockdown system enabled accurate localization of TFP5-3Ty but displayed relatively slow depletion kinetics. In contrast, the mAID system promoted rapid protein degradation with an alteration of the localization. The efficient IAA-induced degradation of TFP5-mAID-HA is consistent with cytosolic accessibility of the C-terminal AID tag, without establishing the overall membrane topology of TFP5. Although the 3Ty tag did not affect parasite fitness, the larger mAID-3HA tag modestly impaired propagation, as reflected by reduced plaque size relative to TIR1 controls. In contrast to the apical localization of TFP5-3Ty, TFP5-mAID-HA accumulated in early secretory compartments, suggesting that the C terminus contributes to proper TFP5 trafficking to the apical pole. Thus, even a partial reduction of apical TFP5 is sufficient to impair egress and invasion, indicating that its function depends on local abundance at the apical pole. Upon complete depletion, both systems showed severe plaque defects. Loss of TFP5 impaired microneme secretion, blocking egress, motility, and invasion, while intracellular replication remained unaffected and no delayed-death phenotype was observed. Despite localizing to compartments involved in secretory organelle trafficking and maturation, TFP5 depletion caused no detectable defects in the trafficking or localization of the rhoptry or microneme proteins. Also, although proteolytic processing of microneme and rhoptry proteins is essential for parasite fitness, TFP5 depletion did not impair the maturation of the proteins examined. Likewise, in TFP5-depleted parasites, Golgi, ELC, and PLVAC markers displayed localization patterns comparable to those observed in control parasites, suggesting that these compartments were not morphologically or functionally affected.

In contrast, TEM revealed striking morphological alterations. First, in the absence of TFP5, the characteristic microneme collar beneath the conoid was lost, resembling the phenotype of the HOOK mutant (57), suggesting that altered membrane lipid composition may impair HOOK-mediated microneme tethering. Second, rhoptries exhibited major morphological defects and failed to cluster efficiently at the parasite apex, indicating that TFP5 contributes to rhoptry maturation and possibly membrane lipid homeostasis rather than early biogenesis.

In this context, structural predictions provide mechanistic insight, revealing strong similarity between TFP5 homologs and the LPC transporter DrMFSD2A, which is proposed to mediate LPC translocation through a “trap-and-flip” mechanism, reorienting LPC molecules across the lipid bilayer (48). Interestingly, TgVMP1, an ER-resident scramblase required for rhoptry morphogenesis, was recently shown to impair microneme secretion upon depletion without affecting microneme morphology or trafficking (58). Unlike TFP5 depletion, TgVMP1 loss also caused intracellular growth defects, likely due to its earlier localization in the secretory pathway, but the overall phenotype remains similar.

Structural analyses raise the possibility that TFP5 transports or reorients LPC, or related lipid species, within intracellular membranes. Through this potential activity, TFP5 may contribute to the organization, positioning, and function of apical secretory organelles. To investigate whether TFP5 depletion affects parasite lipid composition, we performed untargeted lipidomics. Although this approach does not identify transporter substrates directly, it reveals broader lipid alterations associated with TFP5 depletion. These data must be interpreted cautiously, as disruption of intracellular transporters can cause indirect metabolic effects, including lipid accumulation, degradation, or mislocalization. Notably, the TFP5-mAID-HA mutant already exhibits lipid remodeling without IAA treatment, likely reflecting adaptive or compensatory responses to partial TFP5 impairment. Full TFP5 depletion further remodelled the parasite lipidome, with accumulation of DG, oxidized TGs (OxTG), PC, and PE, while SM and hexosylceramides (HexCer) decreased. The accumulation of neutral lipids and oxidized species likely reflect localized perturbations in lipid homeostasis rather than global metabolic dysfunction, as mitochondrial morphology and parasite growth remained unaffected.

PC, PE, and SM are among the most abundant phospholipid classes in rhoptries, with SM enriched relative to the global parasite lipid composition (28). Their dysregulation upon TFP5 depletion may therefore alter rhoptry membrane composition and contribute to the observed morphological defects. Increased phospholipid levels, reduced sphingolipid abundance, and elevated bis(monoacylglycero)phosphate (BMP), a lipid enriched in endolysosomal compartments (59), suggest disrupted intracellular lipid distribution or trafficking. While these data do not identify a specific TFP5 substrate, they support a role for TFP5 in membrane lipid homeostasis.

Collectively, our findings support a model in which TFP5 maintains lipid homeostasis to ensure proper rhoptry morphogenesis and microneme organization and secretion, thereby enabling parasite virulence. These results highlight the importance of the apicomplexan TFPs in apical secretory organelle function and identify them as potential therapeutic targets.

## MATERIAL AND METHODS

### Parasite culture and generation of transgenic lines

#### Parasite culture and transfection

*T. gondii* tachyzoites were maintained in HFFs (ATCC CCD-1112Sk) cultured in DMEM supplemented with 5% FBS, 2 mM L-glutamine and 25 µg/mL gentamicin. RH TIR1 ΔKu80 ΔHXGPRT (60) and DiCre ΔKu80 ΔHXGPRT (61) parasites were used as parental strains to generate the transgenic parasites described in this study. The pSAG1::CAS9-GFP-U6::sgTFP5 plasmid was generated by replacing the sgUPRT sequence in pSAG1::CAS9-GFP-U6::sgUPRT (62). pTub-GRASP55–YFP plasmid (63) was transiently expressed as previously described (64).

### Generation of TFP5-mAID-HA and TFP5-Ty-U1 parasite lines

TFP5-mAID-HA and TFP5-Ty-U1 parasite lines were generated by CRISPR/Cas9-mediated homologous recombination in the TIR1 and DiCre parental strains using repair templates amplified from pYFP-mAID-3HA (65) or pG152-KI-3Ty-lox-SAG1_3’UTR-HX (66), respectively. Transgenic parasites were selected with mycophenolic acid/xanthine and cloned by limiting dilution. Correct integration was verified by PCR, IFA and western blot (**Fig. S2 and Table S2**). Protein depletion was induced with 500 μM IAA or 50 nM rapamycin.

### Immunofluorescence assay

Intracellular tachyzoites grown on HFF monolayers were fixed for 10 min at room temperature with 4% paraformaldehyde and 0.05% glutaraldehyde, quenched with PBS containing 0.1 M glycine, permeabilized with 0.2% Triton X-100 in PBS, and blocked with 3% bovine serum albumin (BSA) in PBS. Primary antibodies (listed in Table S2) were diluted in PBS containing 3% BSA and incubated at room temperature, followed by the appropriate secondary antibodies (Table S2). Coverslips were mounted in Fluoromount-G containing DAPI

### Western Blot

HFF monolayers grown in 6-cm dishes were infected with freshly egressed parasites and incubated for 30 h in the presence or absence of IAA or rapamycin. Infected cells were harvested and lysed directly in SDS-PAGE sample buffer (50 mM Tris-HCl, pH 6.8, 10% glycerol, 2 mM EDTA, 2% SDS, 0.05% bromophenol blue, and 100 mM DTT). Proteins were separated by SDS-PAGE, transferred onto nitrocellulose membranes, and immunoblotted using the primary and HRP-conjugated secondary antibodies listed in Table S2. Membranes were blocked in PBS containing 0.1% Tween-20 and 5% nonfat dry milk, and all antibodies were diluted in the same buffer.

### Solubility assay

TFP5-Ty-U1 parasites were sequentially extracted with PBS, 1 M NaCl, 0.1 M Na₂CO₃, 1% Triton X-100, or 1% SDS. Samples were subjected to five freeze-thaw cycles, incubated on ice for 30 min, and separated into soluble and insoluble fractions by centrifugation. Fractions were analyzed by western blot using anti-Ty antibodies. Catalase, GAP45, and IMC1 served as controls.

### Plaque assay

Confluent HFF monolayers were infected with serial dilutions of freshly egressed *T. gondii* tachyzoites in the presence or absence of IAA or Rapa. After seven days, monolayers were fixed and stained with crystal violet. The lysis plaque areas were quantified by imaging the wells with an EVOS microscope and measuring their areas using ImageJ software.

### Intracellular growth assay

Freshly egressed parasites were allowed to infect confluent HFF monolayers and were cultured for a single round of development (30 hours) or for two successive rounds (48 + 30 hours) in the presence or absence of IAA. The cell cultures were fixed with 4% PFA / 0.05% Glutaraldehyde and processed for IFA using anti-GAP45 antibodies to visualize the parasites. For each condition, the number of parasites per vacuole was quantified by counting 100 vacuoles in three independent biological experiments.

### Egress Assay

Freshly egressed parasites were allowed to invade confluent HFF monolayers and cultured for 30 h in the presence or absence of IAA or for 72 h in the presence or absence of rapamycin. Parasite egress was induced with 10 μM BIPPO for 15 min or DMSO as a control. Cells were fixed with 4% paraformaldehyde/0.05% glutaraldehyde, quenched with PBS containing 0.1 M glycine, and processed for immunofluorescence using anti-GRA3 and anti-GAP45 antibodies to visualize the parasitophorous vacuole and parasites, respectively. At least 100 vacuoles were scored per condition in three independent biological replicates.

### Microneme Secretion Assay

Microneme secretion assays were performed as previously described (46). TFP5-mAID-HA and TFP5-Ty-U1 parasites were treated with IAA for 30 h or Rapa for 72 h, respectively, before stimulation with 2% ethanol or DMSO for 10 min at 37°C. ESA and parasite pellet fractions were prepared and analyzed by Western blot using antibodies against AMA1, MIC2, MIC4, MIC6, catalase, and GRA1. Microneme secretion was quantified by densitometry using ImageJ from three independent biological replicates

### Gliding assay

Syringe-released TFP5-mAID-HA parasites, treated or not with IAA for 30 h, were resuspended in prewarmed DMEM containing 10 μM BIPPO and allowed to glide on 0.1% gelatin-coated coverslips for 15 min at 37°C. Parasites were then fixed with 4% paraformaldehyde/0.05% glutaraldehyde, quenched with PBS containing 0.1 M glycine, and processed for immunofluorescence using anti-SAG1 antibodies to visualize gliding trails.

### Invasion Assay

Syringe-lysed TFP5-mAID-HA and TFP5-Ty-U1 parasites, pretreated or not with IAA for 30 h or Rapa for 72 h, were seeded onto HFF monolayers for 30 minutes before fixation with 4% paraformaldehyde/0.05% glutaraldehyde for 7 minutes. Coverslips were quenched with 1X PBS / 0.1 M glycine for 5 minutes. A first immunodetection was performed using anti-SAG1 antibodies on non-permeabilized cells to label extracellular parasites. Cells were subsequently fixed with 1% formaldehyde in 1X PBS for 7 min, washed with 1X PBS, and permeabilized with 0.2% TX-100 in 1X PBS. After blocking for 20 min with 1X PBS containing 2% BSA, intracellular parasites were stained using anti-GAP45 antibodies. At least 100 parasites were counted per condition in each technical triplicate. The experiment was repeated in three independent biological replicates.

### Transmission electron microscopy (TEM) and FIBSEM

Conventional TEM was performed essentially as previously described (46). HFF monolayers infected with TIR1 or TFP5-mAID-HA parasites were cultured for 30 h in the presence or absence of IAA before fixation, resin embedding, ultrathin sectioning (70 nm), and imaging on a Tecnai 12 G2 transmission electron microscope (FEI) operated at 80 kV. FIBSEM samples were prepared as previously described (40).

### Quantification of rhoptry morphology by transmission electron microscopy

Transmission electron microscopy images of TFP5-mAID-HA parasites cultured in the absence (−IAA) or presence (+IAA) of auxin were analyzed to assess rhoptry morphology. Parasites were classified as displaying either normal or abnormal rhoptry morphology based on their ultrastructural appearance. A total of 43 parasites in the −IAA condition and 36 parasites in the +IAA condition were examined from a single biological experiment. The percentage of parasites exhibiting normal rhoptry morphology was calculated relative to the total number of parasites analyzed for each condition. No inferential statistical analysis was performed, as the quantification was based on a single biological replicate.

### Image acquisition

IFAs were acquired using a Leica TCS SP8 STED microscope equipped with a 63× oil immersion objective (1.4 NA) at the Bioimaging Core Facility of the Faculty of Medicine, University of Geneva.

### Genome mining

The InterPro-predicted MFS Spinster domain of TFP5 was used as a BLASTp query against representative Alveolata genomes available in VEuPathDB (https://veupathdb.org/) and the NCBI non-redundant database (https://blast.ncbi.nlm.nih.gov). Species analyzed and corresponding BlastP scores are provided in **Table S1**.

### Phylogenetic analysis

Phylogenetic analyses were performed on MFS transporters (InterPro: IPR036259) retrieved from UniProt, including orthologous sequences of TgTFP5 from representative Apicomplexan species. Multiple sequence alignment was generated using ClustalW (67) and manually curated to remove misaligned regions, as well as redundant and truncated sequences. The best-fit substitution model was selected using ModelFinder (68) according to the Bayesian Information Criterion. Maximum likelihood phylogenetic reconstruction was performed in IQ-TREE (69) under this model, and node support was evaluated using 1000 bootstrap replicates. The final tree was visualized and annotated using iTOL (70).

### Taxonomy

The taxonomic nomenclature used was obtained from the NCBI Taxonomy database (https://www.ncbi.nlm.nih.gov/taxonomy), accessed in August 2025.

### Structural modeling and binding pocket analysis

TgTFP5 and PfTFP5 3D structures were predicted using AlphaFold3 (71). Structural alignments were performed on PyMOL Molecular Graphics System, Version 3.0 Schrödinger, LLC. Putative binding pockets were identified using ChimeraX (72). Electrostatic potential maps were calculated using the APBS Electrostatics Plugin (73) of PyMOL, with default parameters. Positioning of LPC within DrMFSD2A is based on the published modeling study (74).

### Lipidomic workflow

Untargeted lipidomic analyses were performed essentially as previously described (46). Briefly, confluent HFF monolayers grown in complete DMEM supplemented with 5% FBS were infected with TFP5-mAID-HA parasites in the presence or absence of IAA, or parental TIR1 parasites in the presence of IAA for 48 h. Intracellular parasites were purified by needle disruption and filtration through 3-µm filters, counted, and lipids were extracted using an isopropanol-based protocol containing LPC 18:1-d7 and Cer 18:1-d7/15:0 internal standards. Four technical replicates and three biological replicates per condition were analyzed. Quality control (QC) samples were prepared as previously described (46). Lipid extracts were analyzed by untargeted LC-HRMS using a Vanquish Horizon UHPLC coupled to an Exploris 120 Orbitrap mass spectrometer (Thermo Fisher), using the chromatographic and acquisition parameters described previously (46). Raw data were processed with MS-DIAL v5.5.250221 (75) using the same peak detection, lipid annotation, alignment, and quality-filtering criteria described previously (46). A total of 942 lipid species passed filtering and were retained for downstream analyses. Signal drift across the analytical sequence was corrected using LOESS normalization, followed by probabilistic quotient normalization (PQN) based on median QC intensities to account for biomass differences between samples. PCA and OPLS-DA analyses (Raw and processed Data are detailed in **S3 and S4 Tables)** were performed in SIMCA 17 (Umetrics Sartorius), and statistical analyses were carried out using GraphPad Prism 8. This experimental design allowed us to assess depletion-associated lipidomic changes by directly comparing TFP5-mAID-HA parasites with and without IAA treatment, while using parental TIR1 +IAA parasites as a reference for lipid abundance under auxin-treated conditions. For lipid class comparisons, technical replicates were aggregated within each biological replicate before statistical analysis. Lipid abundances were expressed relative to the parental TIR1 +IAA condition. For the fold-change analysis, the abundance of each lipid class in TFP5-mAID-HA +IAA parasites was divided by that of the corresponding untreated TFP5-mAID-HA replicate from the same biological experiment. Statistical comparisons among the three experimental conditions were performed using a one-way repeated-measures ANOVA followed by Tukey’s multiple comparisons test.

## ACKNOWLEDGMENTS

We thank Dr. M. Lebrun (LPHI), Dr. J.-F. Dubremetz (LPHI), Dr. V. Carruthers (UMMS) and Dr. B. Striepens (UPENN) for providing antibodies. Prof. P. Thompson for providing BIPPO and to J.-B. Marq (UNIGE) for technical assistance. Dr. M. Treeck (GIMM) for the DiCre line; Dr. D. Sibley (WashU) for the TIR1 strain, the provided vectors; Dr. M. Meissner (LMU) for the pG152-KI-3Ty-lox-SAG1_3’UTR-HX construct. We thank the Bioimaging Core Facility team: F. Prodon, O. Brun, and N. Liaudet, for assistance; N. Hulo and our laboratory members for constructive discussions. We acknowledge ToxoDB and VEuPathDB for providing open-access data for the scientific community. This work was funded by grant HUG RC09-18 from the Private Foundation of the Geneva University Hospitals (https://www.fondationhug.org/) and by grant TMAG-3_216166 from the Swiss National Science Foundation (https://www.snf.ch/en), both awarded to D.S.-F. The funders had no role in study design, data collection and interpretation, or the decision to submit the work for publication.

## SUPPLEMENTARY MATERIALS

**Fig. S1.**
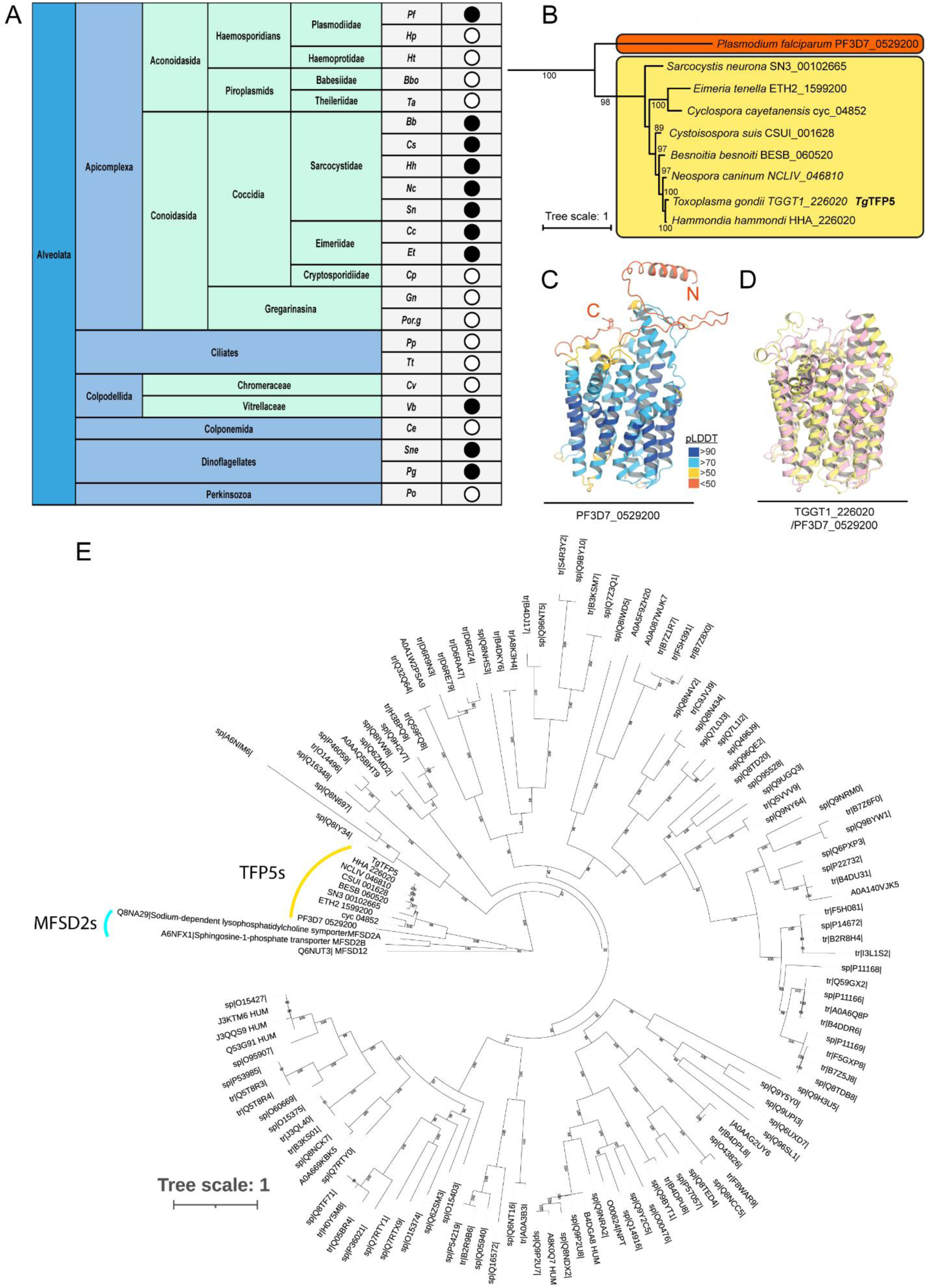
TFP5 is an evolutionarily conserved alveolate transporter related to the human MFSD2 family. **(A)** Black circles indicate species with a conserved TFP5 ortholog, defined using an arbitrary BLASTP *E*-value cut-off of 10⁻²⁰. In contrast, white circles indicate species in which no conserved ortholog was detected. Abbreviations: *Pf*, *Plasmodium falciparum*; *Hp*, *Hepatocystis piliocolobus tephrosceles*; *Ht*, *Haemoproteus tartakovskyi*; *Bbo*, *Babesia bovis*; *Ta*, *Theileria annulata*; *Bb*, *Besnoitia besnoiti*; *Cs*, *Cystoisospora suis*; *Hh*, *Hammondia hammondi*; *Nc*, *Neospora caninum*; *Sn*, *Sarcocystis neurona*; *Cc*, *Cyclospora cayetanensis*; *Et*, *Eimeria tenella*; *Cp*, *Cryptosporidium parvum*; *Gn*, *Gregarina niphandrodes*; *Por.g*, *Porospora gigantea*; *Pp*, *Paramecium primaurelia*; *Tt*, *Tetrahymena thermophila*; *Cv*, *Chromera velia*; *Vb*, *Vitrella brassicaformis*; *Ce*, *Colponema edaphicum*; *Sne*, *Symbiodinium necroappetens*; *Pg*, *Polarella glacialis*; *Po*, *Perkinsus olseni*. **(B)** Phylogenetic reconstruction of TFP5 homologs across the Apicomplexa phylum. The tree was derived from a broader MFS phylogeny of diverse alveolate species and subsequently trimmed to include only Apicomplexan TFP5 sequences. Haemosporidian species are highlighted in orange, whereas coccidian species are shown in yellow. Bootstrap values calculated from 1000 replicates are displayed at the nodes to indicate branch support. The scale bar represents the evolutionary distance, and protein accession numbers are shown for each species. **(C)** AlphaFold3 prediction of the 3D structure of PF3D7_0529200, coloured according to the pLDDT score. The first N-terminal (N) and last C-terminal (C) residues of the predicted protein-coding sequences are annotated on the structure. **(D)** Structural alignment of TgTFP5 (Yellow) and PfTFP5 (Pink) high-confidence regions from AlphaFold models. **(E)** Phylogenetic tree of representative human MFS transporters together with TgTFP5 and conserved apicomplexan homologues (TFP5s). TFP5 homologues form a distinct cluster (indicated in yellow) that groups most closely with the human MFSD2 clade (indicated in cyan). Bootstrap support values are shown at branch nodes, with most internal branches displaying high support. The scale bar represents the number of substitutions per site.

**Fig. S2.**
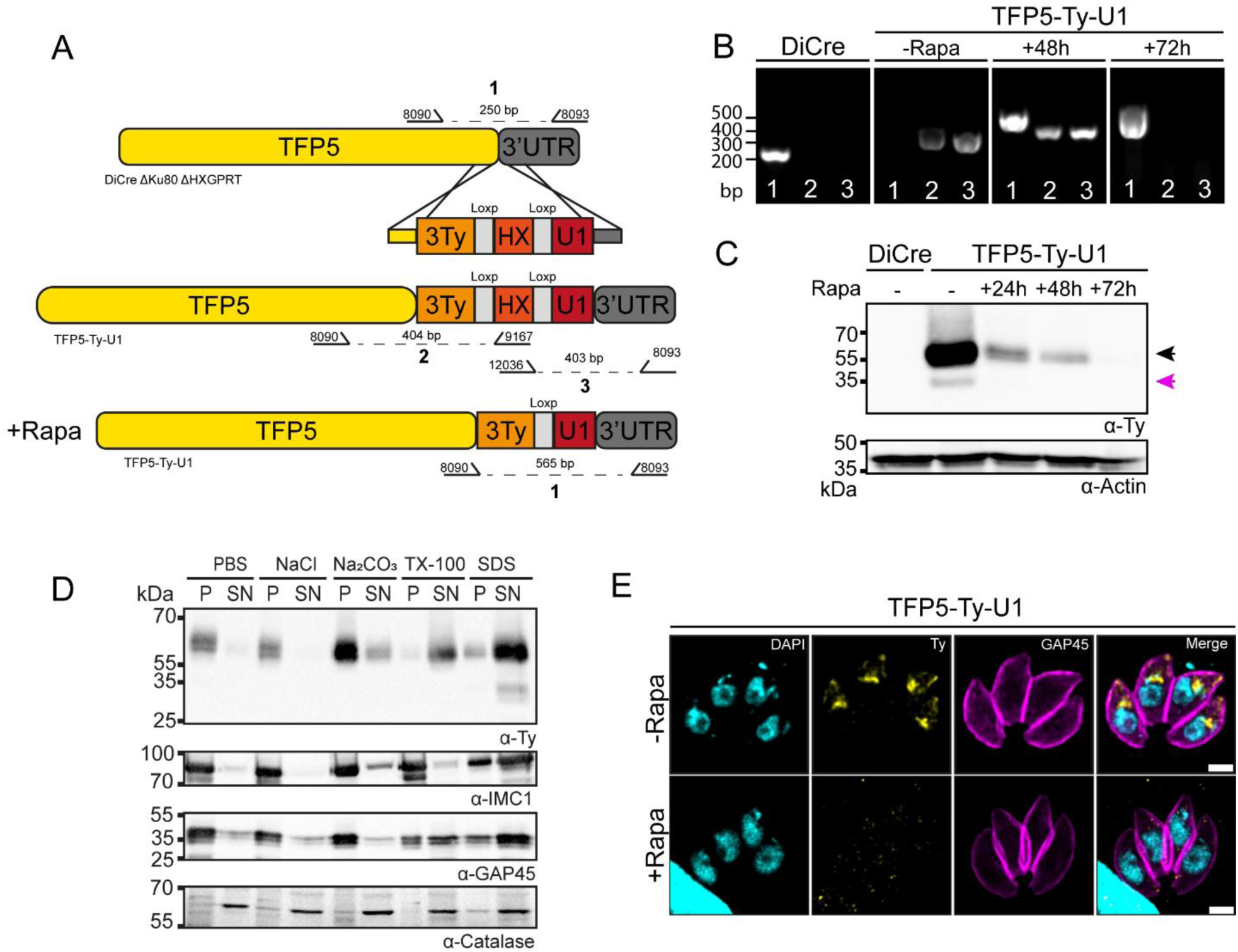
Generation and characterization of the TFP5-Ty-U1 parasite line. **(A)** Schematic overview of the strategy used to generate the TFP5-Ty-U1 parasite line by homologous recombination. Primer pairs used to identify the WT and modified loci following Rapa treatment (1), as well as those detecting the insertion of the 3Ty-HX-U1 cassette (2 and 3), are indicated by black arrows. **(B)** PCR analysis performed on DiCre and TFP5-Ty-U1 parasites after various durations of Rapa treatment, revealing fragments corresponding to the WT, recombined, and excised versions of the TFP5 locus. **(C)** Western blot analysis monitoring TFP5 depletion in intracellular parasites. Parental DiCre and TFP5-Ty-U1 strains were either treated with Rapa or left untreated. TFP5 was detected with an anti-Ty antibody, while anti-Catalase was used as a loading control. The full-length protein is marked by a black arrow, whereas additional tagged protein forms are indicated by purple arrows. **(D)** Solubility analysis of TFP5-3Ty supporting its prediction as a polytopic membrane protein. Parasites were disrupted by freeze–thaw cycles and subsequently incubated with different buffers (PBS, 1 M NaCl, 0.1 M Na₂CO₃, 1% TX-100, 1% SDS). Samples were then separated into supernatant (SN) and pellet (P) fractions. Detection of TFP5-3Ty was carried out using anti-Ty antibodies. Anti-IMC1 and anti-GAP45 antibodies were used as controls for TX-100 insoluble proteins and TX-100 soluble proteins. Anti-Catalase served as a marker for soluble proteins. **(E)** Immunofluorescence analysis of intracellular TFP5-Ty-U1 parasites treated or not with Rapa, 72 hours post-infection, demonstrating the loss of TFP5 signal (anti-Ty; yellow). The IMC was visualized using anti-GAP45 (magenta), and nuclei were stained with DAPI (cyan). Scale bar: 2 µm.

**Fig. S3.**
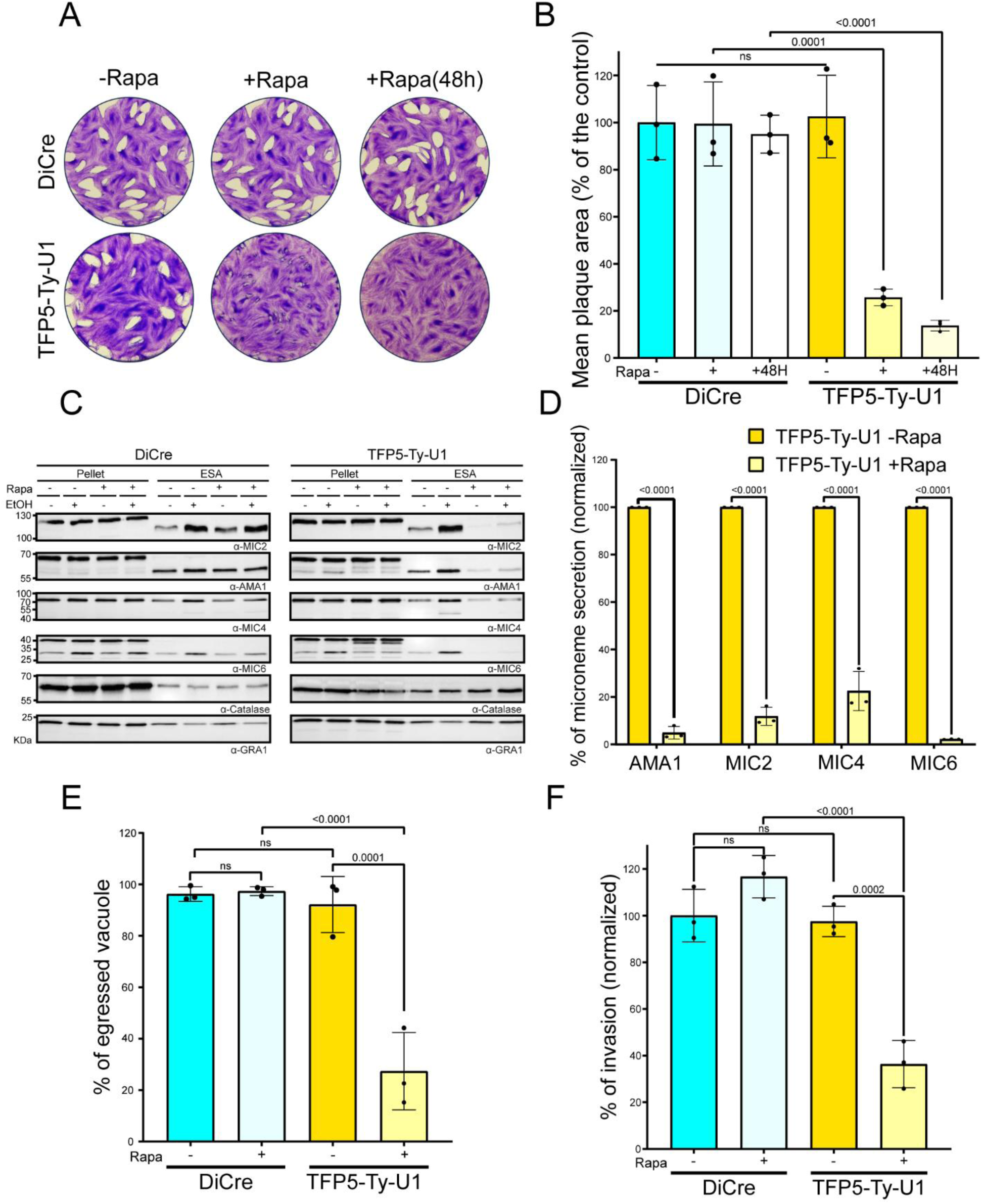
Effects of Rapamycin-induced TFP5 depletion on parasite microneme secretion, egress and invasion. **(A)** Representative images from plaque assays conducted across three independent experiments illustrating lytic cycle progression in parental DiCre and TFP5-Ty-U1 parasites under three conditions: without Rapa treatment, with Rapa treatment, or after 48 h of Rapa pretreatment before infection. **(B)** Quantification of plaque assay outcomes for DiCre and TFP5-Ty-U1 parasites in the presence or absence of Rapa. Plaque areas were normalized relative to the DiCre –Rapa condition. Statistical significance was evaluated using a one-way ANOVA followed by Tukey’s multiple comparison test (mean ± SD; n = 3 independent biological replicates). **(C)** Western Blot analysis of microneme secretion in DiCre and TFP5-Ty-U1 parasites treated with or without Rapa following ethanol (EtOH) stimulation. Secretion of MIC2, AMA1, MIC4, and MIC6 was evaluated by examining their presence in pellet (P) and supernatant (S) fractions. Anti-GRA1 served as a control for constitutive secretion, while catalase was used to monitor parasite lysis. Shown are representative images from three independent biological replicates. **(D)** Quantification of AMA1, MIC2, MIC4, and MIC6 released into the supernatant after EtOH stimulation. Secretion levels were normalized to constitutive GRA1 secretion for each sample and then expressed relative to the TFP5-Ty-U1-Rapa condition. Statistical analysis was performed using two-way ANOVA followed by Tukey’s multiple comparisons test (mean ± SD; n = 3 independent biological replicates). **(E)** Quantification of parasite egress in DiCre and TFP5-Ty-U1 parasites with or without Rapa treatment after stimulation with the egress-inducing compound BIPPO. Data were analyzed for statistical significance using one-way ANOVA followed by Tukey’s multiple comparisons test (mean ± SD; n = 3 independent biological replicates). **(F)** Invasion assay of intracellular DiCre and TFP5-Ty-U1 parasites following 72 h of pretreatment with or without Rapa. Invasion efficiency was measured 30 minutes post-infection and normalized to the DiCre-Rapa control. Statistical significance was evaluated using one-way ANOVA with Tukey’s multiple comparisons test (mean ± SD; n = 3 independent biological replicates).

**Fig. S4.**
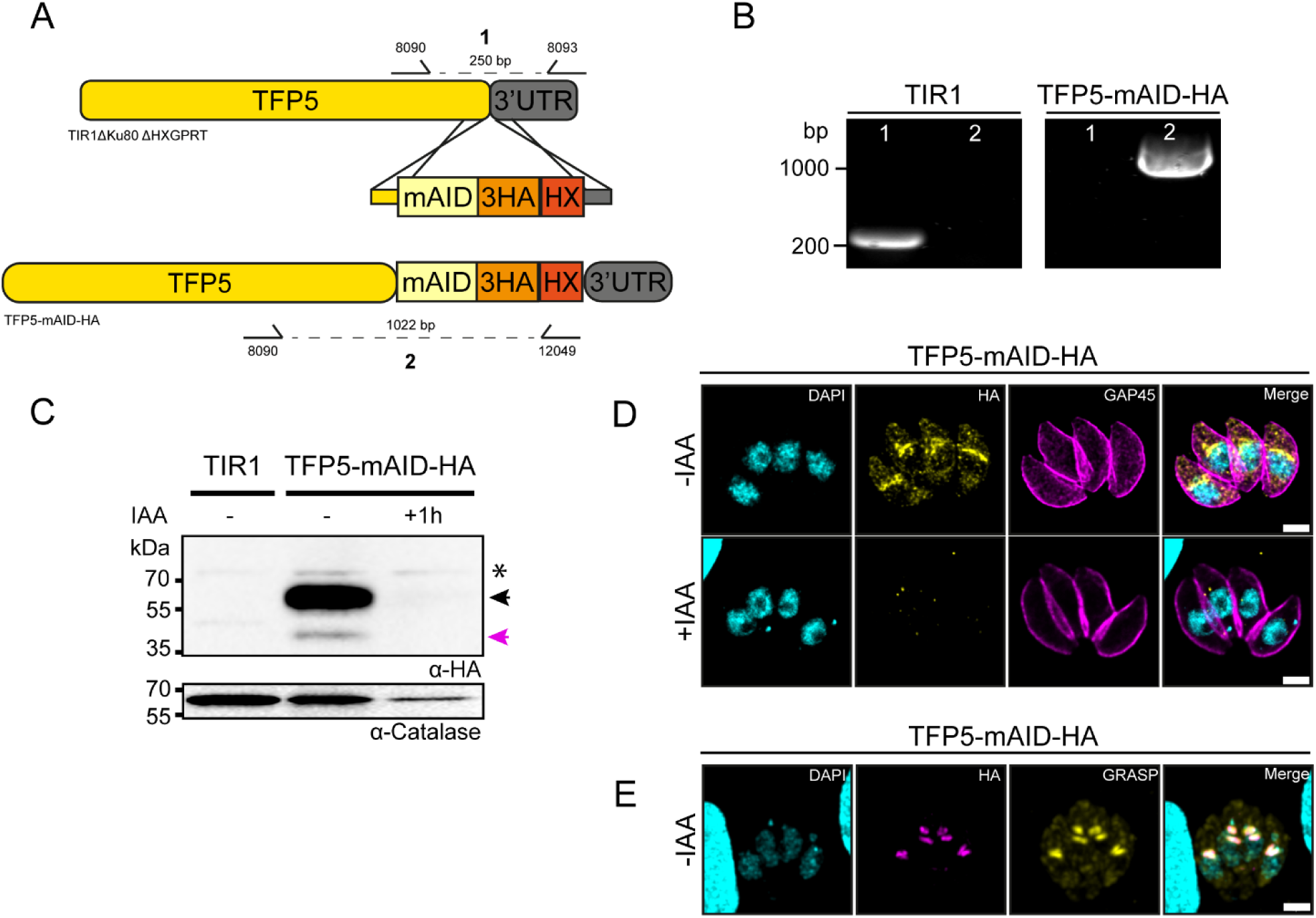
Generation, validation, and subcellular localization of the TFP5-mAID-HA mutant. **(A)** Scheme illustrating the generation of the TFP5-mAID-HA parasite line through homologous recombination. Primer pairs designed to amplify the WT locus (1) and the integrated mAID-3HA-HX cassette (2) are shown by black arrows. **(B)** Agarose gel electrophoresis displaying PCR products corresponding to either the WT or recombined TFP5 locus in TIR1 and TFP5-mAID-HA strains. **(C)** Immunoblot analysis showing the degradation kinetics of the TFP5-mAID-HA-3HA protein. Parental TIR1 and TFP5-mAID-HA parasites were incubated with IAA or left untreated. Detection of TFP5 was performed using an anti-HA antibody, with anti-Catalase used as a loading control. The full-length protein is indicated by a black arrow, and additional tagged forms are highlighted with purple arrows. An asterisk denotes a non-specific signal. **(D)** IFA of intracellular TFP5-mAID-HA parasites ± IAA treatment, 30 hours post-infection, showing the disappearance of the TFP5 signal (anti-HA; yellow). The IMC was labeled with anti-GAP45 (magenta), and DNA was stained with DAPI (cyan). **(E)** IFA of TFP5-mAID-HA parasites showing colocalization of TFP5-mAID-HA (magenta) with the Golgi marker GRASP (yellow). Nuclei were stained with DAPI (cyan). Scale bar: 2 µm.

**Fig. S5.**
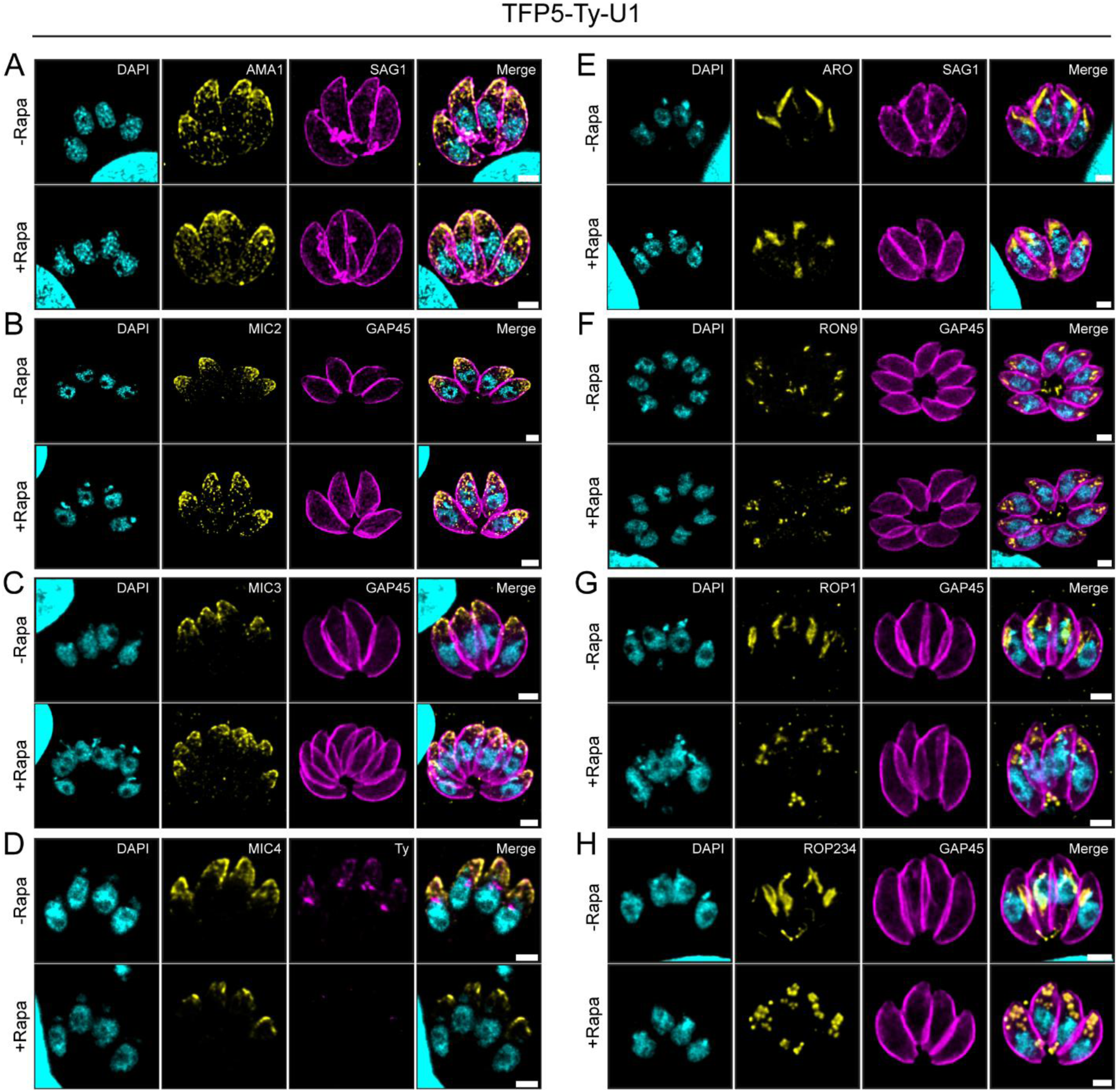
TFP5 depletion preserves microneme trafficking while altering rhoptry morphology in both U1 and mAID systems. **(A–D)** Microneme organization is maintained upon TFP5-Ty-U1 iKD. Parasites were stained with **(A)** anti-AMA1, **(B)** anti-MIC2, **(C)** anti-MIC3, or **(D)** anti-MIC4 (yellow), together with anti-SAG1 to label the plasma membrane, anti-GAP45 to visualize the IMC or anti-Ty to label the TFP5-3Ty protein (magenta). **(E–H)** Rhoptry morphology is disrupted following TFP5 depletion. Parasites were stained with **(E)** anti-ARO, **(F)** anti-RON9, **(G)** anti-ROP1, or **(H)** anti-ROP2-4 (yellow), with anti-GAP45 or SAG1 (magenta), respectively marking the IMC and plasma membrane. Nuclei were labeled with DAPI in all images. Scale bars, 2 µm.

**Fig. S6.**
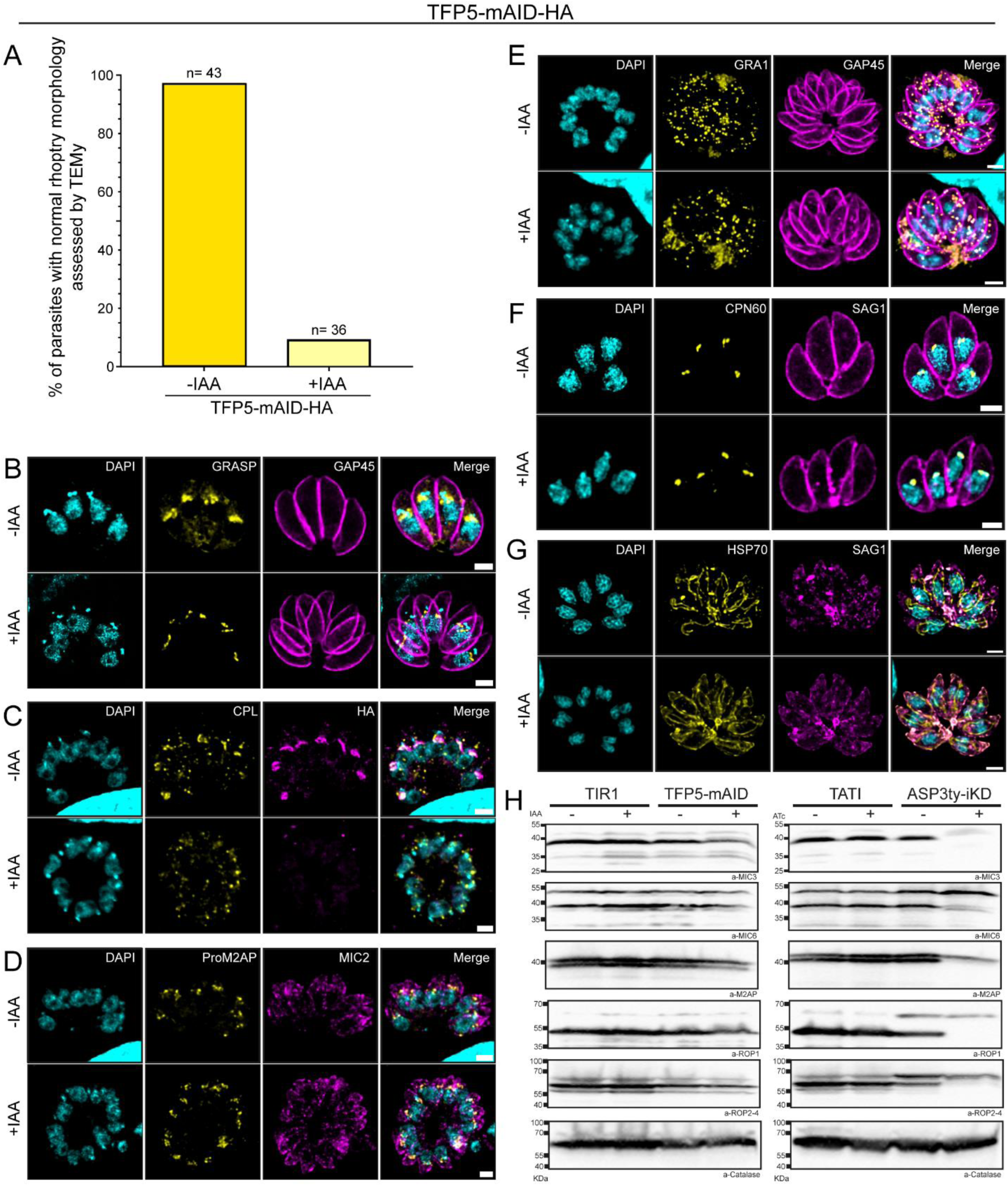
TFP5 depletion preserves Golgi and post-Golgi organelle integrity and does not impair apically secreted protein maturation. **(A)** Quantification of rhoptry morphology following TFP5 depletion. Percentage of TFP5mAID parasites displaying normal rhoptry morphology in the absence (−IAA) or presence (+IAA) of auxin, as assessed by transmission electron microscopy. A total of 43 parasites were examined under control conditions (−IAA) and 36 parasites following auxin treatment (+IAA). Quantification was performed from a single biological experiment. Data are presented as descriptive percentages without statistical analysis. (**B–G)** TFP5 depletion doesn’t impact the staining pattern of the following organelles. IFA showing **(B)** the cis-Golgi using GRASP55-YFP transient expression (yellow), **(C)** the PLVAC using the CPL marker (yellow) and **(D)** the ELC, using proM2AP (yellow). The integrity of the dense granules (**E**), apicoplast (**F**), and mitochondria (**G**) was assessed using anti-GRA1, anti-CPN60, and anti-HSP70 antibodies, respectively (yellow). Anti-GAP45 or SAG1 has been used to delineate parasite shape, anti-MIC2 to label the micronemes and anti-HA has been used to mark the TFP5-mAID-HA protein (magenta). DAPI (cyan) stains nuclei. Scale bars: 2 µm. **(H)** Depletion of TFP5 does not impair the proteolytic maturation of microneme and rhoptry proteins, in contrast to ASP3 depletion. Western blot analysis of apical secretory protein maturation in TIR1, TFP5-mAID, TATI, and ASP3ty-iKD parasite lines grown in the absence (-) or presence (+) of IAA or ATc, as indicated. Blots were probed for MIC3, MIC6, M2AP, ROP1, and ROP2–4. Catalase was used as a loading control.

**Fig. S7.**
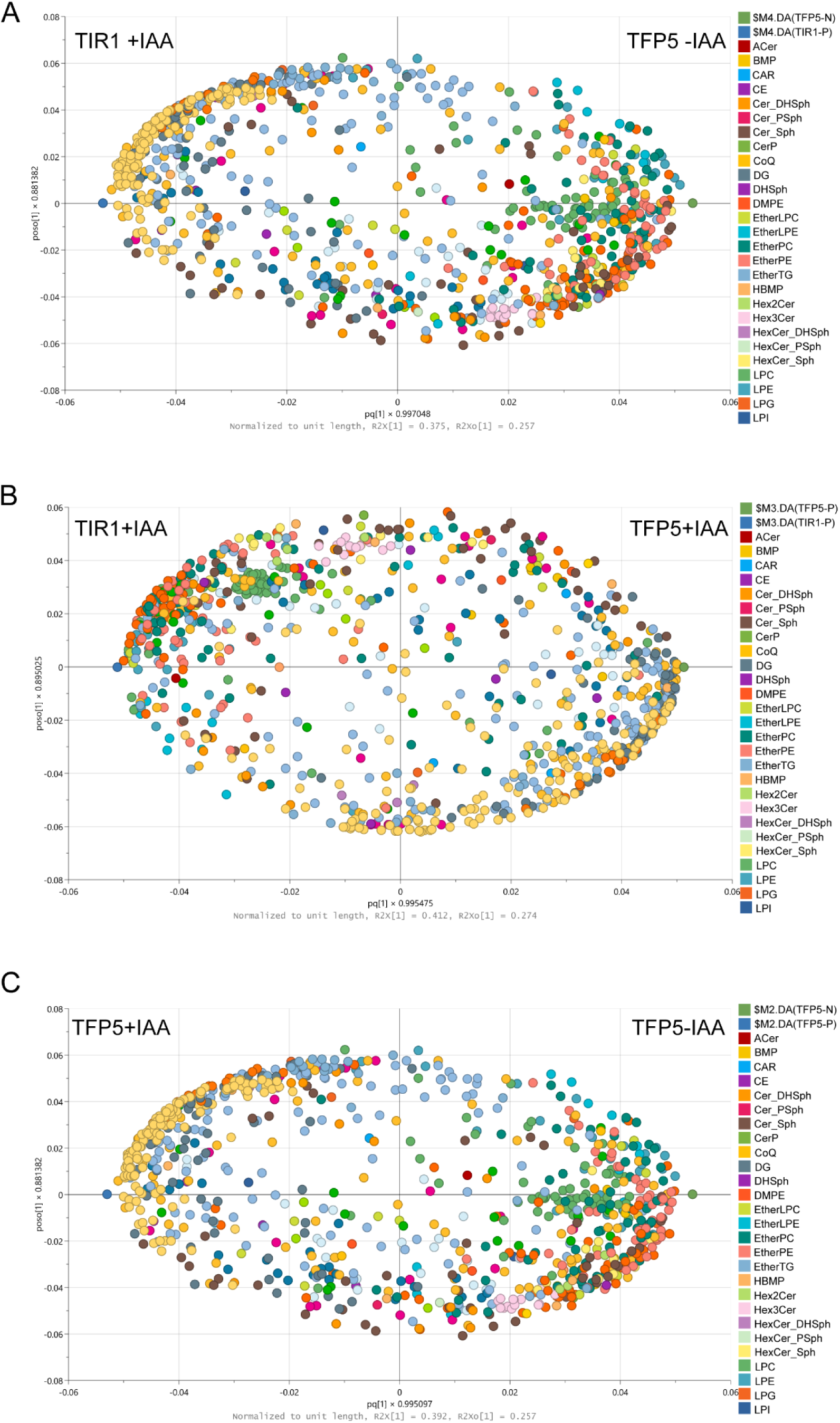
OPLS-DA identifies lipid signatures associated with TFP5 depletion. (**A–C**) Score plots showing the loadings derived from OPLS-DA models comparing (**A**) TIR1 +IAA versus TFP5-mAID-HA-IAA (1 + 3 + 0; R²X = 0.84, Q² = 0.98), (**B**) TIR1 +IAA versus TFP5-mAID-HA +IAA (1 + 1 + 0; R²X = 0.69, Q² = 0.97), and (**C**) TFP5-mAID-HA −IAA versus TFP5-mAID-HA +IAA (1 + 1 + 0; R²X = 0.65, Q² = 0.92). The predictive component (pq(1)) represents the variation associated with the experimental conditions, whereas the orthogonal component (poso(1)) captures variability independent of group separation. Each dot represents an individual lipid species, color-coded according to its lipid class, and its position along the x-axis reflects its contribution to group discrimination. For instance, in panel A, lipids with positive pq(1) values are more abundant in TFP5-mAID-HA-IAA, while those with negative values are enriched in the TIR1 +IAA condition. This representation allows the identification of lipid classes that drive the separation between conditions. TFP5-mAID-HA −IAA was used as the baseline to assess lipidomic changes associated with TFP5 depletion, whereas TIR1 +IAA served as a control for nonspecific effects of IAA treatment.

**Fig. S8.**
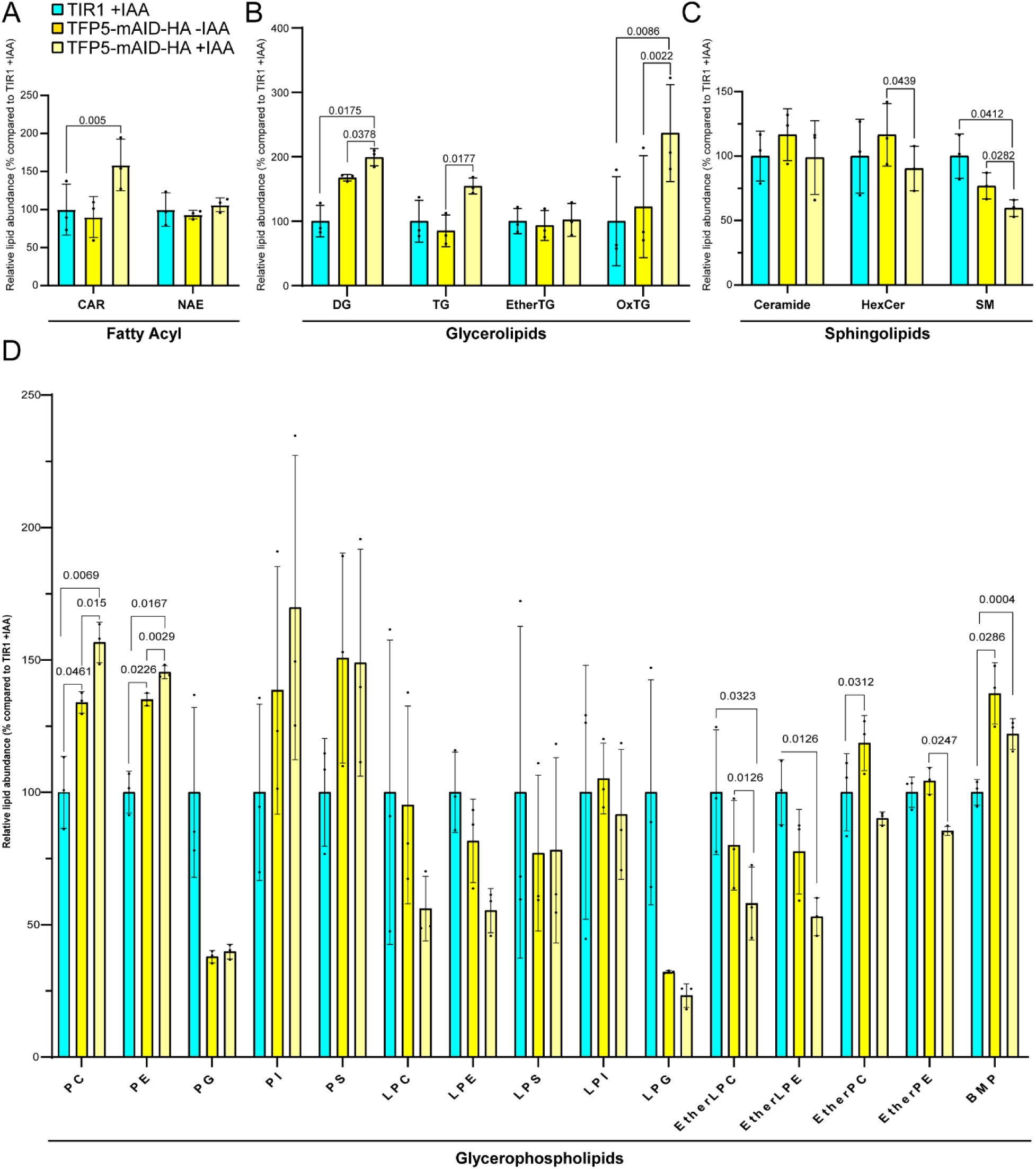
TFP5 depletion alters the lipid composition of *T. gondii* parasites. Global lipidomic profiling was performed on control parasites (TIR1 +IAA, cyan) and TFP5-mAID-HA parasites cultured in the absence (-IAA, yellow) or presence (+IAA, light yellow) of auxin. Lipid abundance is presented relative to the TIR1 +IAA condition. **(A)** Analysis of fatty acyl-derived lipids, including acylcarnitines (CAR) and N-acylethanolamines (NAE). **(B)** Quantification of neutral glycerolipids, including diacylglycerols (DG), triacylglycerols (TG), ether-linked TGs (EtherTG), and oxidized TGs (OxTG). **(C)** Analysis of sphingolipid classes, including ceramides, hexosylceramides (HexCer), and sphingomyelins (SM). **(D)** Quantification of glycerophospholipid species, including phosphatidylcholine (PC), phosphatidylethanolamine (PE), phosphatidylglycerol (PG), phosphatidylinositol (PI), phosphatidylserine (PS), lysophospholipids (LPC, LPE, LPS, LPI, LPG), ether phospholipids (EtherLPC, EtherLPE, EtherPC, EtherPE), and bis(monoacylglycero)phosphate (BMP). Data are presented as mean ± SD with three individual biological replicates shown as black dots. All three experimental conditions were analyzed in parallel within each biological experiment. Statistical significance was determined using one-way ANOVA followed by Tukey’s multiple comparisons test. Adjusted p-values are indicated on the graphs.

